# Glutamine availability regulates cDC subsets in tissue

**DOI:** 10.1101/2024.09.17.613574

**Authors:** Graham P. Lobel, Nanumi Han, William A. Molina Arocho, Michal Silber, Jason Shoush, Michael C. Noji, Tsun Ki Jerrick To, Li Zhai, Nicholas P. Lesner, M. Celeste Simon, Malay Haldar

**Affiliations:** Department of Pathology and Laboratory Medicine, Perelman School of Medicine, University of Pennsylvania, Philadelphia, PA, 19014, USA; Abramson Family Cancer Research Institute, Perelman School of Medicine, University of Pennsylvania, Philadelphia, PA, 19104, USA; Department of Cancer Biology, Perelman School of Medicine, University of Pennsylvania, Philadelphia, PA, 19014, USA; Department of Cell and Developmental Biology, Perelman School of Medicine, University of Pennsylvania, Philadelphia, PA, 19014, USA; Institute for Immunology, Perelman School of Medicine, University of Pennsylvania, Philadelphia, PA, 19104, USA

## Abstract

Proliferating tumor cells take up glutamine for anabolic processes engendering glutamine deficiency in the tumor microenvironment. How this might impact immune cells is not well understood. Using multiple mouse models of soft tissue sarcomas, glutamine antagonists, as well as genetic and pharmacological inhibition of glutamine utilization, we found that the number and frequency of conventional dendritic cells (cDC) is dependent on microenvironmental glutamine levels. cDCs comprise two distinct subsets – cDC1 and cDC2, with the former subset playing a critical role in antigen cross-presentation and tumor immunity. While both subsets show dependence on Glutamine, cDC1s are particularly sensitive. Notably, glutamine antagonism did not reduce the frequency of DC precursors but decreased proliferation and survival of cDC1s. Further studies suggest a role of the nutrient sensing mTOR signaling pathway in this process. Taken together, these findings uncover glutamine dependence of cDC1s that is coopted by tumors to escape immune responses.

**One Sentence Summary:** Type 1 conventional dendritic cells require glutamine to maintain their number in non-lymphoid tissue.

**Significance:** Immune evasion is a key hallmark of cancer; however, the underlying pathways are diverse, tumor-specific and not fully elucidated. Many tumor cells avidly import glutamine to support their anabolic needs, creating a glutamine-deficient tumor microenvironment (TME). Herein, using mouse models of soft tissue sarcomas, we show that glutamine depletion in TME leads to reduced type 1 conventional dendritic cells – a cell type that is critical for adaptive immune responses. This work is a paradigm for how tumor cell metabolism can regulate anti-tumor immune responses and will be foundational to future efforts targeting glutamine metabolism for cancer immunotherapy.

## INTRODUCTION

In malignant or other highly proliferative cells, glutamine is a “conditionally essential” amino acid due to its involvement in a wide variety of anabolic cellular processes (1). Glutamine contributes to tricarboxylic acid (TCA) cycle anaplerosis, *de novo* nucleotide and non-essential amino acid synthesis, control of reactive oxygen species via glutathione production , protein and lipid glycosylation, and signaling through mechanistic target of rapamycin complex 1 (mTORC1) (1). Due to its multifaceted role in supporting cell growth and proliferation, glutamine metabolism has long represented an attractive target for anti-cancer therapy (2). The first glutamine related compound tested was 6-diazo-5-oxo-L-norleucine (DON), a pan-glutamine antagonist that blocks all glutaminergic pathways, but toxicity limited its clinical effectiveness (2). Over the past decade, work in this area has focused on inhibiting glutaminolysis, the conversion of glutamine to glutamate via the mitochondrial enzyme glutaminase (GLS), which cancer cells become “addicted to” to fuel anabolic processes (1). CB-839, a selective GLS1 inhibitor, has demonstrated efficay against some *in vivo* tumor models and is currently being tested in multiple clinical trials in combination with a variety of other anti-cancer drugs (3–8). However, other *in vivo* tumor models have shown that CB-839 is ineffective, and clinical trial results thus far have demonstrated mixed results (8–10).

One explanation put forth to explain these variable outcomes is that tumor cells efficiently rewire their glutamine metabolic pathways to avoid strict dependence on GLS, despite its central role in glutamine catabolism. To avoid this possibility, recent work has been directed at “rediscovering” DON by synthesizing prodrugs that are only activated upon entry into the tumor microenvironment. One such compound, JHU-083, has demonstrated robust responses in multiple *in vivo* models of breast, melanoma, and colorectal cancer (11–13). Most strikingly, JHU-083 has also demonstrated impressive synergy with immune checkpoint blockade therapy (13). This finding is surprising, because glutamine metabolism is a critical regulator of T cell differentiation and effector functions (14–18). Counterintuitively, however, several groups using a variety of glutamine-blocking compounds have repeatedly demonstrated that the inhibition of glutamine uptake or metabolism does not negatively affect intratumoral T cell function and instead synergizes with immune checkpoint blockade (19–21).

In contrast to T cells, the *in vivo* effects of glutamine blockade on intratumoral myeloid cell functions are less well-understood and deserving of further study, as myeloid cells make up the vast majority of tumor resident immune cells (22). Inhibition of glutamine metabolism can suppress the production of myeloid-derived suppressor cells (MDSCs) in both humans and mice, and may additionally increase MHCII expression in murine macrophages (12, 21). How glutamine metabolism regulates other critical intratumoral myeloid cells remains unclear.

Dendritic cells (DCs) are professional antigen-presenting cells that constantly surveil lymphoid and non-lymphoid organs for potential threats, migrate into lymph nodes upon activation, and initiate T cell responses when needed (23). DCs can be divided by function and ontogeny into several subsets with type 1 conventional DCs (cDC1s) especially important for anti-tumor immune responses (24). These subsets arise from pre-specified hematopoietic precursors, which, in mice, circulate in the form of pre-cDC1s and pre-cDC2s, and terminally differentiate upon organ entry, where they also upregulate organ-specific transcriptional programs and surface markers (25). The relative frequency of cDC1s vs. cDC2s differs between tissues but cDC1 frequency is particularly low in solid tumors (24). Indeed, many studies have shown that the frequency (based on overall cDCs) and numbers of cDC1s correlate positively with T cell infiltration and responses to immunotherapy (26–30). Thus, the cellular and molecular basis of low cDC1s in tumors represent an important knowledge gap with therapeutic implications.

We find here that glutamine availability differentially affects cDC1 and cDC2 frequency in tumors and normal tissue. cDC1s are particularly sensitive to microenvironmental glutamine – a phenomenon that is also recapitulated in DCs generated *in vitro* using established FLT3 ligand-based culture systems. Using genetic and pharmacological approaches *in vivo* and *in vitro*, we provide evidence that a combination of reduced survival, proliferation, and maturation of cDC1s underlie its sensitivity to low glutamine. Our findings reveal how tissue glutamine levels can impact antigen presentation through regulation of cDC1s, which has important implications for immune responses. These insights will also help inform ongoing efforts to develop glutamine antagonism for cancer therapy.

## RESULTS

### Glutamine differentially regulates cDC subsets in the tumor microenvironment

In order to assess the importance of glutamine in tumor-associated DCs, we employed the pan-glutamine antagonist DON (**Figure S1A**). Treatment of an implantable methylcholanthrene-induced model of fibrosarcoma (FS tumors) with DON inhibited tumor growth without overt toxicity based on mouse body weight (**Figures 1A and S1B**). To examine DON’s effects on intratumoral myeloid cells, we quantified populations of macrophages, monocytic MDSCs (M-MDSCs), polymorphonuclear MDSCs (P-MDSCs), and DCs in DON and vehicle-treated tumors via flow cytometry (**Figure S1C**). In accord with previous work, reduced M-MDSC-like cells were observed with DON administration (**Figure S1C**). Notably, DCs were also significantly decreased in DON-treated tumors (**Figure 1B**). Further examination showed that cDC1 frequency (as a percentage of cDCs) and absolute numbers were particularly diminished upon DON treatment (**Figure 1C**). Due to its relatively greater influence on cDC1s, DON exposure increased the frequency (as a percentage of cDCs) of cDC2s while still reducing their overall numbers within tumors (**Figure 1C**).

**Figure 1:**
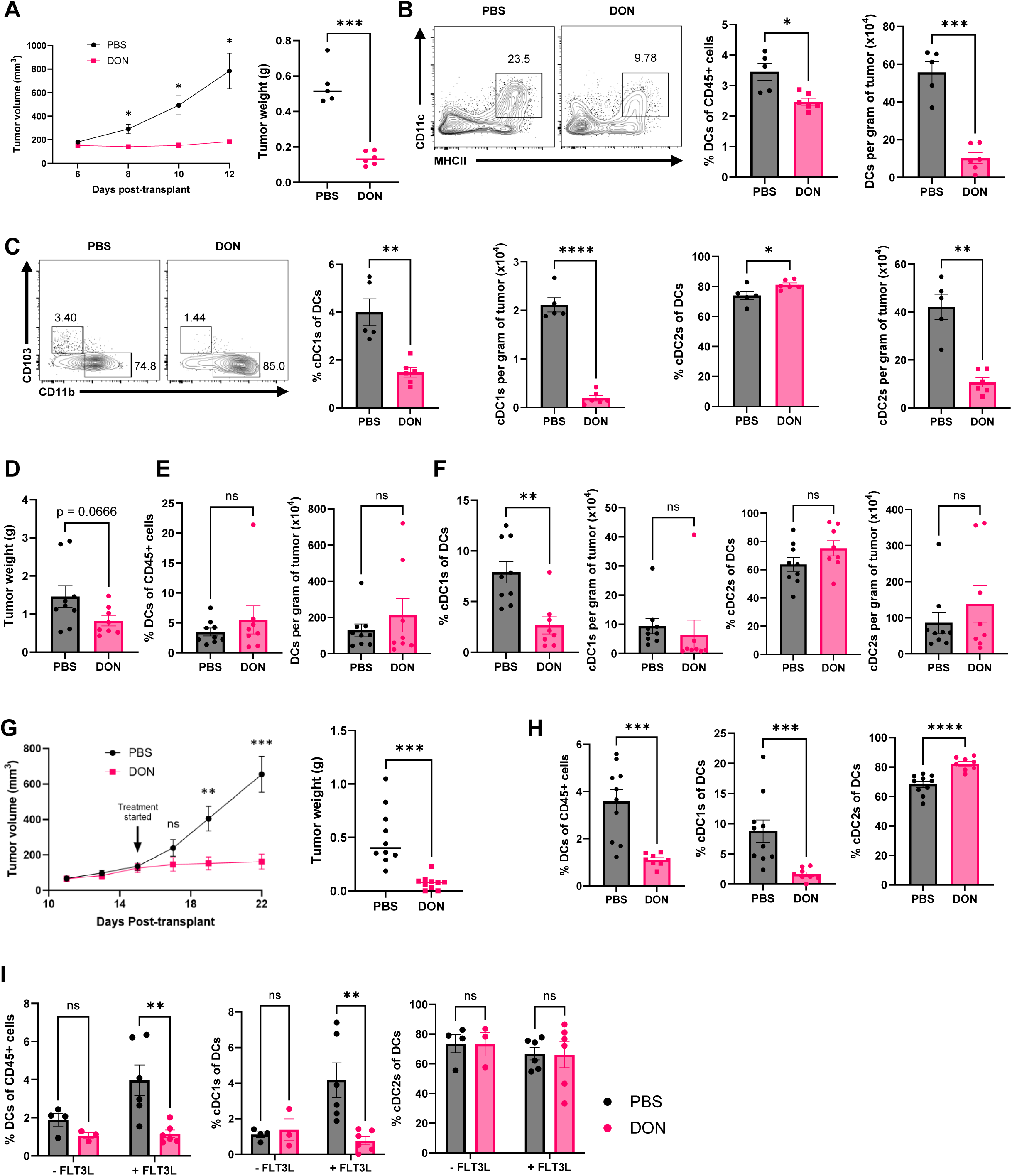
Glutamine antagonism reduce CDCs in the tumor microenvironment A: Volumes (left) and weights (right) of subcutaneous syngeneic FS tumors treated daily (intraperitoneal, starting after six days post tumor cell transplantation) with PBS or 0.25 mg/kg DON. Tumor volumes were measured every two days and weighed at endpoint. Experiment repeated ≥ 3 time. B: Flow cytometry-based identification (left) and quantification (right) of intratumoral DCs analyzed at the endpoint of the experiment described in 1A. DCs were defined as CD45^+^ Ly6c^-^ Ly6g^-^ F4/80^-^ CD11c^+^ MHCII^+^ cells. C: Flow cytometry-based identification (left) and quantification (bar graphs) intratumoral DC subsets at the endpoint of the experiment described in 1A. cDC1s were defined as CD103^+^ CD11b^-^ DCs, while cDC2s were defined as CD103^-^ CD11b^+^ DCs. D: Weights of tumors in a genetically engineered mouse model of undifferentiated pleomorphic sarcoma (UPS) treated (intraperitoneal injections) daily for 12 days with PBS or 0.25 mg/kg DON. Treatments were started after tumors were detected and weights were quantified by subtracting the mass of the contralateral leg from that of the tumor-bearing leg. Experiment repeated ≥ 3 time. E-F: Bar graphs quantifying intratumoral DCs (as defined as in 1B) as well as the two major DC subsets (as defined in 1C) at the endpoint of experiment described in 1D. G: Tumor volumes and weights from transplantable syngeneic UPS-bearing mice treated with either PBS or 0.5 mg/kg DON for 7 days. H: Bar graphs quantifying the proportions of intratumoral DCs among CD45^+^ cells, and cDC1s and cDC2s within DCs, in syngeneic transplantable UPS tumors after 7 days of treatment with PBS or 0.5 mg/kg DON. I: Bar graphs quantifying the proportions of intratumoral DCs among CD45^+^ cells, and cDC1s and cDC2s within DCs, in FS tumors cells after 6 days of treatment with PBS or 0.25 mg/kg DON +/- FLT3L injections.

We next validated these findings in an independent autochthonous mouse model of undifferentiated pleomorphic sarcoma (UPS), generated by injecting TAT-Cre protein into the hindlimb musculature of *LSLKras^G12D/+^;Trp53^fl/fl^* mice (31). Animals were treated with DON once tumors were detected and no changes in overall mouse weight were observed (**Figure S1D**). Although DON treatment showed a trend towards reduced tumor mass, the difference did not reach statistical significance due to large variability in tumor sizes (**Figures 1D**). However, cDC1 frequency was significantly reduced by DON without impacting overall cDC frequency (**Figures 1E-F**). We also tested this in a syngeneic implantable UPS model by treating with 0.5 mg/kg DON – finding strong effects on tumor growth (**Figure 1G**), no decrease in mouse weights (**Figure S1E**), and significant depletion of intratumoral DCs and cDC1s in particular (**Figure 1H**).

Other studies have shown that cDC1 numbers can be increased in tumors by injection of FLT3 ligand (FLT3L) – a key cytokine in DC-poiesis produced by several tumor resident immune cells (29, 30). As such, we tested whether DON acts by reducing FLT3 ligand levels in the tumor microenvironment. FS-bearing mice were treated with PBS or DON in the presence or absence of FLT3L (**Figure 1I**). Notably, the FLT3L-induced boost in intratumoral DCs and cDC1s was suppressed by DON treatment, further underscoring the importance of glutamine to cDC1 infiltration in this context.

Amongst tumors, STS are particularly avid users of glutamine, mostly converting it into glutamate through GLS for further catabolism (6). In fact, implantable and spontaneous STS models were particularly sensitive to GLS inhibition in comparison to other *in vivo* tumor models (10). Importantly, GLS1 deletion in tumor cells has previously been shown to increase glutamine levels in tumor interstitial fluid (19). Therefore, *shRNAs* to deplete GLS1 in our FS cell line were employed in an attempt to increase the availability of glutamine to intratumoral immune cells (**Figures 2A-B**). As anticipated, GLS1 Knockdown (GLS-KD) reduced glutamine uptake by FS tumor cells from the surrounding media (**Figure 2C**). We then implanted GLS-KD cells into syngeneic mice. In contrast to complete *Gls1* gene deletion or total GLS blockade through DON, the knockdown approach preserves some GLS activity in tumor cells allowing them to grow *in vivo* (**Figures 2D - E**). Notably, GLS-KD tumors harbored more DCs and cDC1s compared to control (*shScramble*) FS tumors (**Figure 2F – G**). Taken together, these results underscore the importance of glutamine in regulating cDC1 numbers and frequency in solid tumors.

**Figure 2:**
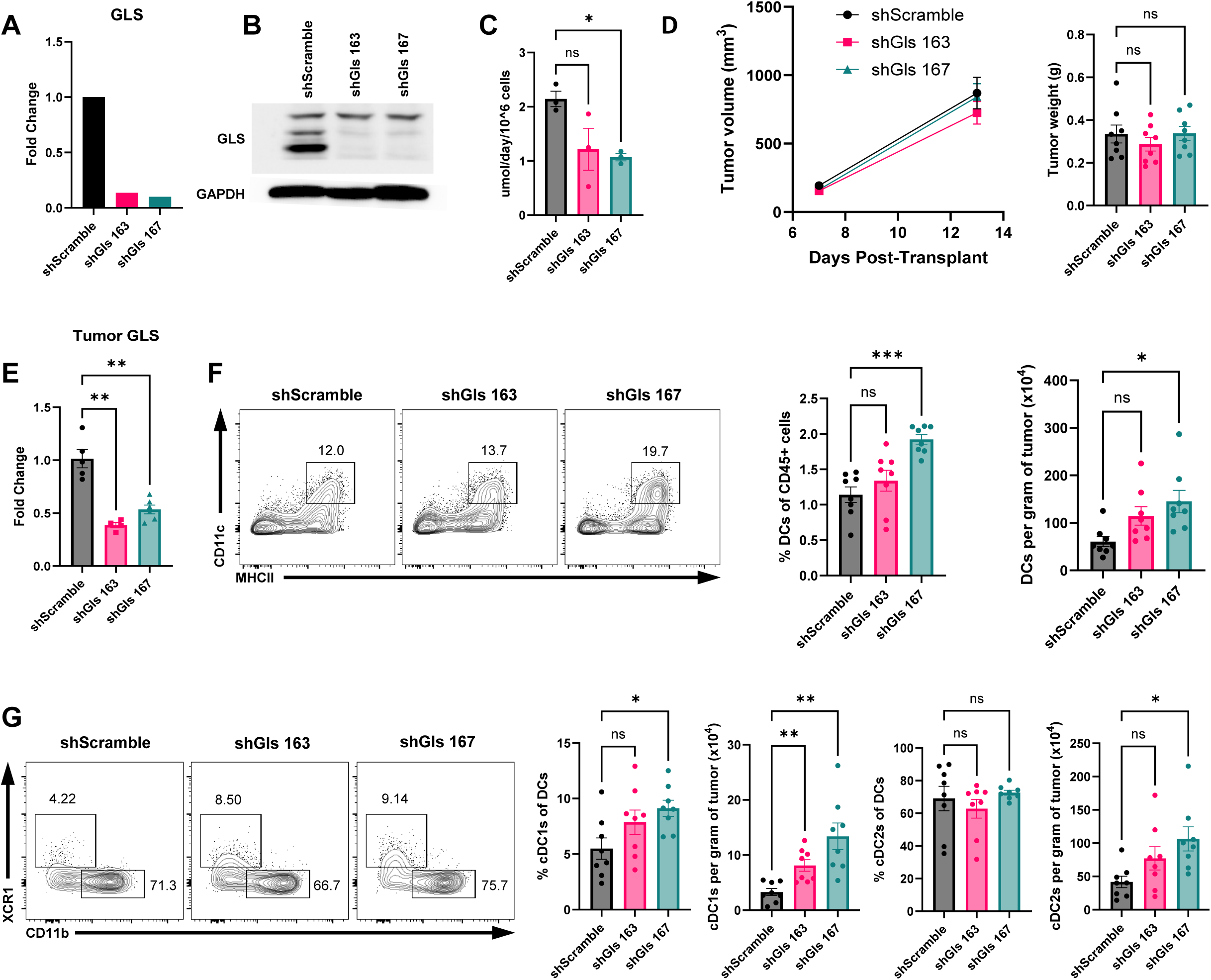
Glutamine levels regulate CDC distribution in the tumor microenvironment A: Glutaminase (*Gls*) transcript levels (relative to *Hprt* and normalized to shScrampble expressing cells) in FS cell lines expressing shScramble (control) and two different GLS-targeting short hairpins as measured by RT- qPCR. B: Western blot showing expression of GLS protein in control and GLS knockdown cells described in 2A. C: Quantification of glutamine uptake per cell in control and GLS knockdown cells described in 2A. Uptake was calculated by quantifying glutamine concentration (using an YSI metabolite analyzer) in tumor-conditioned media compared to cell-free control after 24 hours, then normalized to cell number. D: Volumes (left) and weights at endpoint (right) of shScramble and shGLS tumors (syngeneic flank transplantation into C57BL/6 mice). T cells were depleted every three days to avoid confounding effects of potential anti-tumor T cell responses. Experiment repeated ≥ 3 times. E: Expression of *Gls* transcripts (relative to *Hprt* and normalized to control shScramble tumors) in shScramble *vs.* shGLS knockdown tumors quantified by RT-qPCR. F: Flow cytometry-based identification (left, flow cytometry plots) and quantification (bar graphs) of total cDCs (as defined in Figure 1B) as a percentage of CD45^+^ cells (left bar graph) as well as per gram of tumor (right bar graph) in control and GLS-knockdown tumors described in 2D. G: Flow cytometry-based identification (left, flow cytometry plots) and quantification (bar graphs) of cDC1s, (XCR1^+^ CD11b^-^ DCs) and cDC2s (XCR1^-^ CD11b^+^ DCs) as a percentage of all DCs as well as per gram of tumor in control and GLS-knockdown tumors described in 2D.

### Glutamine regulates cDC subsets in normal tissue

While the above findings demonstrate a glutamine dependence for tumor microenvironmental cDC1s, it was unclear whether the same holds true in normal tissues. To investigate this, we treated non-tumor-bearing WT mice with vehicle or DON for 10 days and examined DC distribution in different organs. Similar to the tumor microenvironment, we found that DCs, and especially cDC1s, were depleted from lung (**Figures 3A-B**), liver (**Figures S2A - B**), and inguinal lymph nodes (**Figure S2C**). To further ensure that these effects were related to glutamine blockade and not non-specific effects of DON, we treated mice with V9302 – an inhibitor of SLC1A5, a major glutamine transporter - for 10 days and found outcomes consistent with DON treatment (Figures 3C-D).

**Figure 3:**
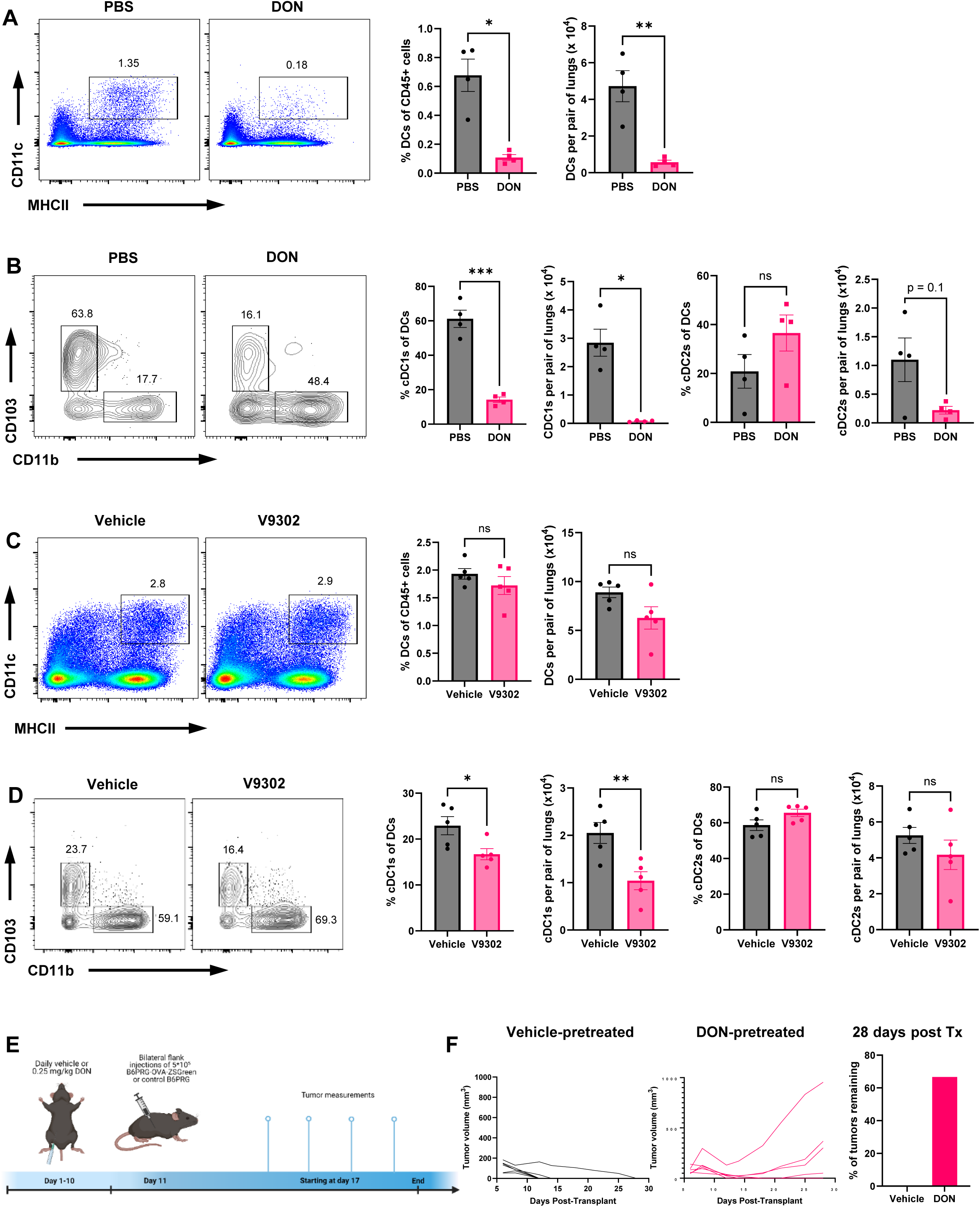
Glutamine regulates tissue CDC distribution A: Representative flow cytometric plots (left) and quantification (bar graphs, right) of lung DCs (CD45^+^ Ly6c^-^ Ly6g^-^ CD64^-^ CD11c^+^ MHCII^+^ cells) after PBS or DON treatment. Bar graphs show indicated cell type as a percentage of CD45^+^ cells (left graph) and total number per pair of lungs (right graph). B: Representative flow cytometric plots (left) and bar graphs (right) quantifying lung cDC1s and cDC2s, as defined in Figure 1C, after PBS or DON treatment. Bar graphs show indicated cell type as a percentage of DCs or total number of cells per pair of lungs. C: Representative flow cytometric plots (left) and bar graphs (right) quantifying total lung DCs (as defined in Figure 3A), after vehicle or V9302 treatment. Bar graph show indicated cell type as a percentage of CD45^+^ cells or total cells per pair of lungs. D: Representative flow cytometric plots (left) and bar graphs (right) quantifying lung cDC1s and cDC2s (as defined in Figure 1C), after vehicle or V9302 treatment. Bar graphs show indicated cell type as a percentage of DCs or total number of cells per pair of lungs. E: Experimental schematic of DON pretreatment and OVA-ZSGreen fibrosarcoma transplantation experiment. F: Tumor volume curves for OVA-ZSGreen fibrosarcoma growth in vehicle- and DON-pretreated and quantification of the proportion of tumors remaining 28 days after transplantation.

Next, we examined functional consequences of reduced tissue cDC1 with glutamine antagonism, using anti- tumor immune responses as a functional readout. Briefly, mice were pretreated with DON for 10 days to deplete/reduce cDC1s and then transplanted with a modified version of FS tumor cells engineered to express the model antigen OVA and the ZSGreen reporter and then monitored for tumor growth (**Figure 3E**). As anticipated, PBS-treated mice fully rejected OVA-containing tumors. In contrast, DON-treated mice were not able to fully reject these highly immunogenic tumors (**Figure 3F**). The outgrowth of tumors in DON pretreated mice is particularly striking given that DON also suppresses tumors in a tumor-cell intrinsic manner. We concluded that cDC1 deficiency induced by DON treatment leads to defective sensing of transplanted immunogenic tumors leading to reduced adapative immune responses. Overall, these findings highlight the requirement of glutamine for normal cDC1 numbers and functions.

### Glutamine supports differentiation, proliferation and survival of cDCs

The differential effects of DON on cDC1s prompted an investigation into the cellular basis of these observations. Specifically, we first determined whether glutamine antagonism suppresses DC development and subsequent differentiation. In mice, DC precursors committed to becoming either cDC1s or cDC2s (pre-cDC1s and pre-cDC2s, respectively) are generated in the bone marrow from macrophage-DC progenitors through common DC precursors. Pre-cDCs then circulate and terminally differentiate after organ entry (**Figure 4A**) (25). To assess whether glutamine blockade negatively impacts DC precursor production, we treated mice with DON for 6 days, then harvested lung and bone marrow and quantified DC precursors and subsets. While mature cDC populations in the lungs were significantly depleted by DON (**Figure S3A**), pre-DCs and pre-DC1s in the bone marrow were not affected (**Figures 4B-C and S3B**). Correspondingly, circulating pre-DCs identified using *Zbtb46-GFP* mice were also unaffected with 10 days of DON treatment (**Figure S3C**). Zbtb46 is a DC-specific transcription factor and replacement of the coding sequence by an EGFP reporter molecule previously established a mouse line (*Zbtb46-GFP*) that faithfully identifies the DC lineage (32). These results suggest that the effect of DON on tissue DC subsets is unlikely due to effects on precursor specification.

**Figure 4:**
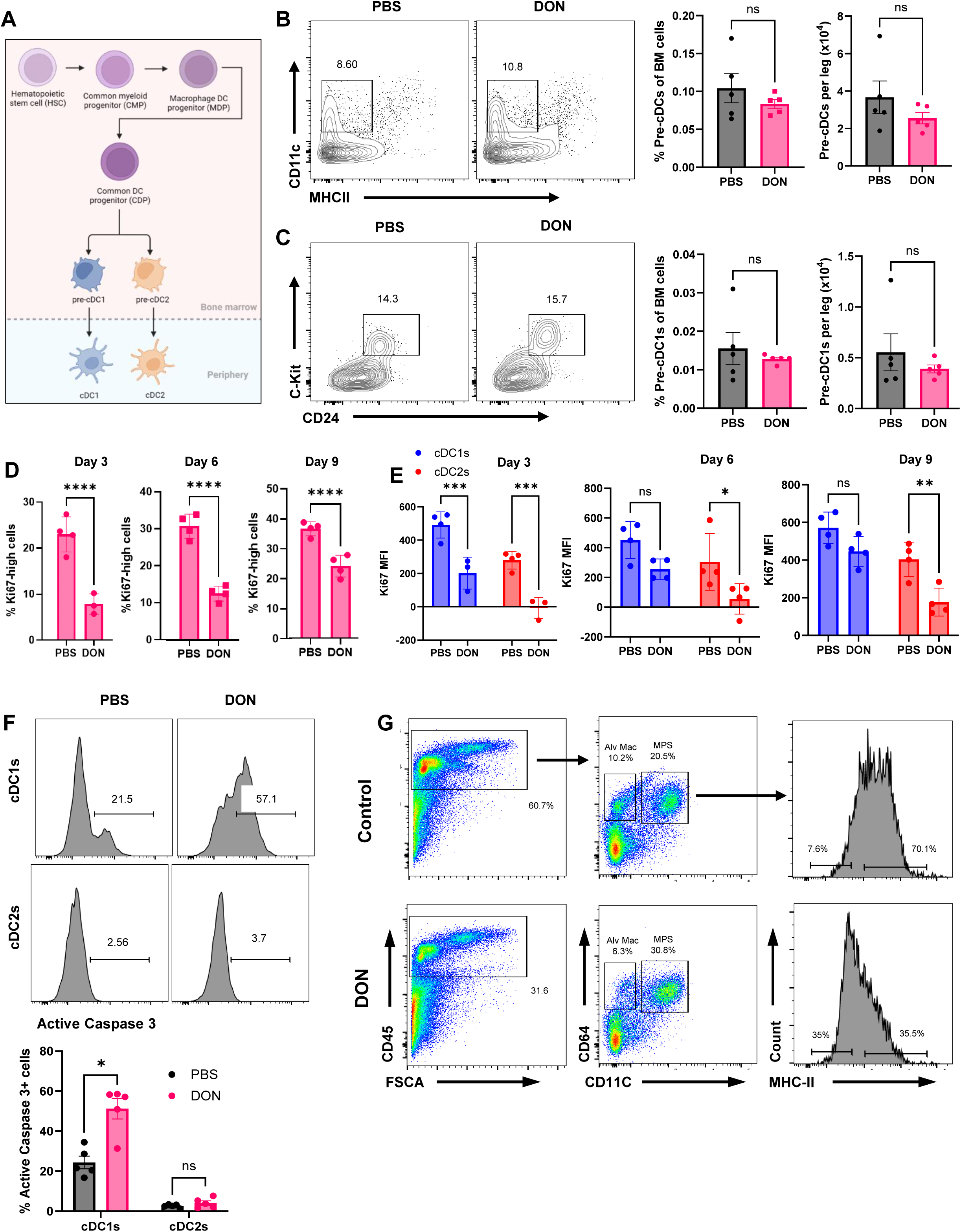
Glutamine differentially regulates CDC subsets in vivo A: Diagram illustrating DC development B: Representative flow cytometric plots and bar graphs quantifying pre-DCs, defined as Lin^-^ SiglecH^-^ Flt3^+^ CD11c^+^ MHCII ^low^ cells, after PBS or DON treatment as a percentage of bone marrow cells and per leg. C: Representative flow cytometric plots and bar graphs quantifying pre-DC1s, defined as CD117^+^ CD24^+^ pre- cDCs, after PBS or DON treatment as a percentage of bone marrow cells and per leg. D: Bar graphs quantifying the proportion of Ki67^+^ cells among lung DCs via flow cytometry after PBS or DON treatment for the specified duration (header). E: Bar graphs quantifying the expression of Ki67 among lung cDC1s and cDC2s via flow cytometry after PBS or DON treatment for the specified duration (header). F: Representative histograms and bar graphs quantifying the proportion of Active Caspase 3^+^ cells among lung cDC1s and cDC2s after 6 days of PBS or DON treatment. G: Lungs from mice treated (IP) with PBS or DON (0.5 mg/kg) for 7 days were subjected to flow cytometric analyses for the frequency of myeloid antigen presenting cells.

Differentiated tissue DCs display limited proliferation (33–35). Thus, we next examined whether reduced cDC1 proliferation can explain its response to glutamine antagonism by treating mice with DON for 3, 6, and 9 days and examining DC proliferation through Ki67 staining (**Figures 4D-E**). Reduced proliferation of lung DCs was apparent within three days of treatment. This reduction, however, was not specific to cDC1s and also observed in cDC2s precluding cell proliferation as the primary driver of the selective effect on cDC1s. Thus, we next investigated whether glutamine antagonism differentially affects survival of DC subsets using active Caspase 3 staining (**Figure 4F**). Significantly increased active Caspase-3 levels were detected in lung cDC1s compared to cDC2s. Of note, we noticed that DON treatment led to increased CD11C^high^ MHCII^low^ cells consistent with immature or precursor DCs **(Figure 4G**). Therefore, increased cCDC1 cell death alongside a block in pre-cDC1 terminal differentiation could explain a differential impact on cDC subsets, while reduced proliferation of both CDC1 and CDC2 accounts for the overall reduction in cDCs upon glutamine antagonism.

### Glutamine levels correlate with cDC1 differentiation

Aforementioned *in vivo* studies engendered profound glutamine deficiency through DON treatment. As technical limitations prevent precise alterations of glutamine levels *in vivo*, we adopted a well-established *in vitro* culture system of DC-poiesis that produces a mixture of plasmacytoid DCs, cDC1s, and cDC2s. Consistent with our *in vivo* studies, we observed that overall DCs, and especially cDC1s, were lower upon reducing media glutamine concentrations in a dose dependent manner **(Figure 5A-B**). At lower levels, the generation of cDC1s was selectively impaired relative to cDC2s (**Figure 5B**). We further validated these findings in an unbiased manner through single-cell RNAseq (sc-RNAseq), which showed a greater impact of low glutamine or DON exposure on cDC1 frequency **(Figure 5C**). Of note, we identified a population of cells expressing generic DC lineage markers such as Zbtb46 but low levels of subset-specific or maturation markers (**Figure S4A-B**). We therefore labelled them as cDC precursors (pre-cDC) as we could not categorize them under known DC subsets (**Figures 5C and S4A-B**). Importantly, this pre-DC-like population expanded under glutamine restrictive conditions, which is consistent with our *in vivo* observations above regarding a differentiation block from pre-CDC to cDC1.

**Figure 5:**
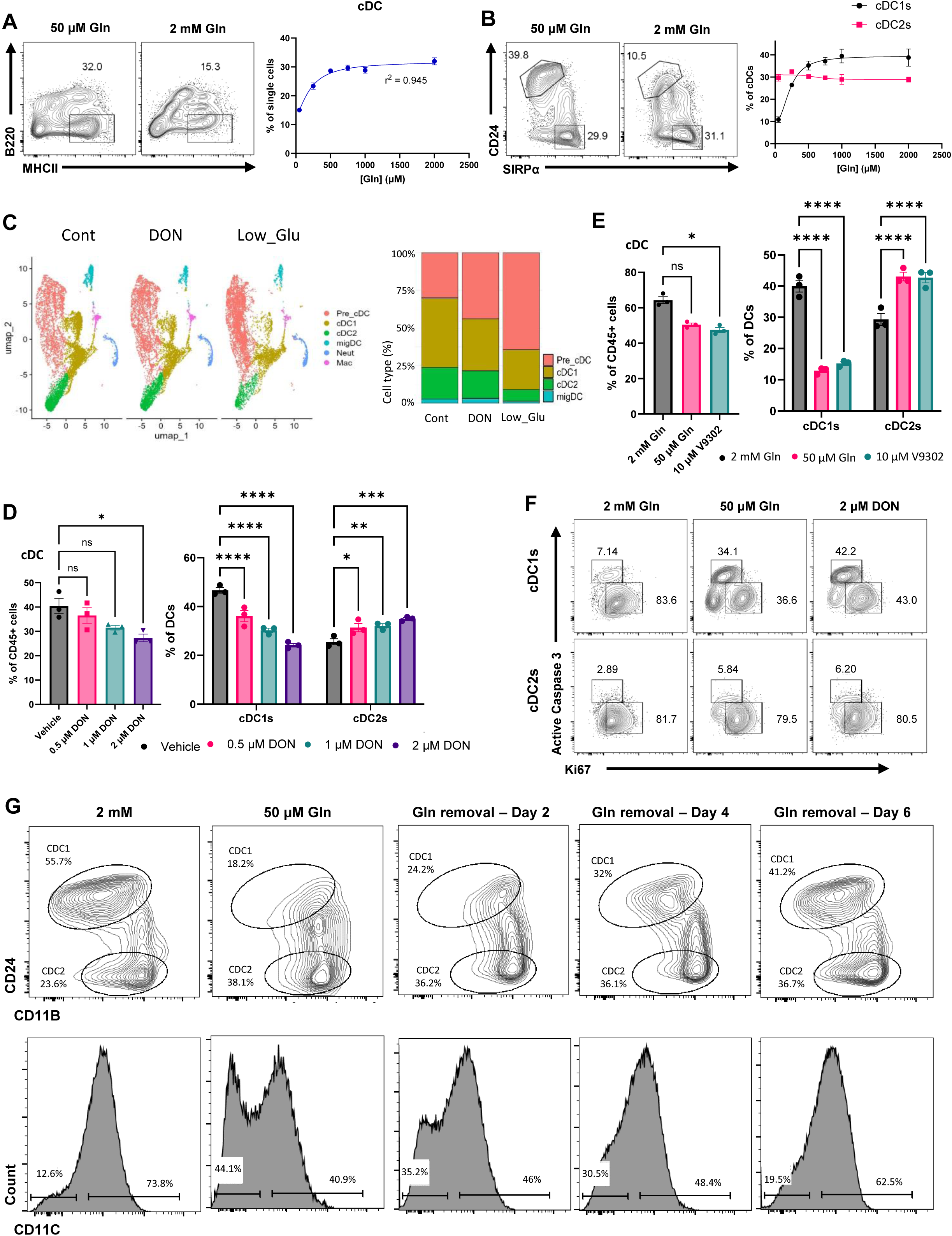
Glutamine differentially regulates CDC subsets in vitro A: Representative flow cytometric plots and dose-response curve quantifying the proportion of FLT3L-derived BMDCs, defined as B220^-^ MHCII^+^ cells after 7 days of culture with varying concentrations of glutamine. B: Representative flow cytometric plots and dose-response curves quantifying the proportions of FLT3L- derived cDC1s, defined as CD24^+^ SIRPα^-^ DCs, and cDC2s, defined as CD24^-^ SIRPα^+^ DCs, among DCs after 7 days of culture with varying concentrations of glutamine. C: UMAP plots on the left shows distribution of cDC subsets in scRNA-seq of FLT3L-derived BMDCs after 7 days of culture with regular glutamine (2000 µm), low glutamine (50 µm), or DON. The UMAP shows only cells that express *Zbtb46*, as the selected marker for DCs. *Zbtb46* negative cells were discarded from the analyses. cDC1, cDC2, and migratory DCs (mDCs) were identified by expression of characteristic markers of these cell types (see supplementary Figure S4B). Minor contamination with neutrophils and macrophages were also identified. The bar graph on right plots the percentage (Y-axis) of each cDC subset (color-coded bars) under D: Bar graphs quantifying the proportion of FLT3L-derived BMDCs (leftmost panel), defined as CD45^+^ B220^-^ MHCII^+^ cells, among CD45^+^ cells, and cDC1s and cDC2s (middle and right panels) among cDCs via flow cytometry after culture with the specified doses of DON for 9 days. E: Bar graphs quantifying the proportion of FLT3L-derived BMDCs (leftmost panel), among CD45^+^ cells, and cDC1s and cDC2s (middle and right panels) among cDCs via flow cytometry after culture with either 2 mM Gln, 50 μM Gln, or 2 mM Gln with 10 μM V9302 for 9 days. F: Representative flow cytometric plots showing proportions of Ki67^+^ Active Caspase 3^-^ proliferating cells and Ki67^-^ Active Caspase 3^+^ apoptotic cells among cDC1s and cDC2s after culture for 5 days with either 2 mM Gln, 50 μM Gln, or 2 mM Gln with 2 μM DON. Please see supplementary Figure S5A for the corresponding bar graph from quantification. G: Representative flow cytometric plot showing overall DCs and the Cdc subsets in FLT3 cultured bone marrow derived DCs that were cultured under normal conditions and glutamine levels for 5 days, after which glutamine deficiency were induced. Cells were analyzed 2, 4, 6 days after glutamine manipulation.

Similar to altering glutamine levels, DON exposure showed dose-dependent ability to reduce cDC1 frequency (**Figure 5D**). Opposing glutamine uptake with V9302 also reproduced the effects of glutamine antagonism on cDC1 frequency (**Figure 5E**). Given these overall similarities between *in vivo* and *in vitro* effects of glutamine on cDC subsets, we next tested whether glutamine differentially regulates proliferation and survival of *in vitro* generated DC subsets (**Figure 5F**). cDC1 proliferation was selectively impaired under low glutamine conditions, in contrast to *in vivo* findings where DON treatment affected both cDC1 and cDC2 proliferation equally. However, consistent with our *in vivo* findings, low glutamine levels or DON treatment induced significantly more apoptosis in cDC1s compared to cDC2s (**Figure 5F**). To disentangle the impact of glutamine on DC survival/proliferation from its effects on pre-cDC differentiation, we cultured BMDCs under normal glutamine conditions and then imposed glutamine deficiency at day 2, day 4, and day 6 followed by flow cytometric analyses at day 8. Importantly, the impact of glutamine on cDC1 frequency diminished with time (**Figure 5G**). Furthermore, there was less accumulation of pre-cDC-like CD11C^low^ MHCII^+^ cells when glutamine deficiency was administered at later time points (**Figure 5G**). Overall, low glutamine appears to affect differentiation, proliferation, and survival of cDC1s.

### Glutamine regulates CDC subsets through mTORC1 signaling

Glutamine is utilized by a variety of metabolic and signaling pathways to support cell growth and proliferation (**Figure 6A**), where one or several of these contribute to the regulation of overall cDC and cDC1 numbers. To address this, we systematically queried the role of key metabolic pathways downstream of glutamine, starting with glutaminolysis as the conversion of glutamine to glutamate via its rate-limiting enzyme - mitochondrial GLS - represents a significant source of biosynthetic metabolites to support cell proliferation (36). We treated differentiating BMDCs with the selective GLS inhibitor CB-839, which did not significantly impact the production of total cDCs or cDC1s (**Figure 6B**). Alpha-ketoglutarate (αKG) is a key metabolite of glutaminolysis that contributes to TCA cycle anaplerosis as well as TET-mediated DNA demethylation (37). αKG can also be converted to glutamate via transaminases and elevate multiple metabolites generated by glutamine catabolism (38). Thus, we assessed whether restoring flux through this pathway rescues cDC and cDC1 production in the absence of glutamine by adding cell-permeable αKG to glutamine-deprived media **(Figure 6C**), as well as to media containing DON (**Figure S5B**). Remarkably, while αKG restored overall cDCs, it failed to rescue the selective depletion of cDC1s. This result is somewhat counterintuitive, given that GLS inhibition had little effect on cDC or cDC1 generation, implying that mitochondrial GLS may not be the main producer of glutamate or αKG during cDC differentiation.

**Figure 6:**
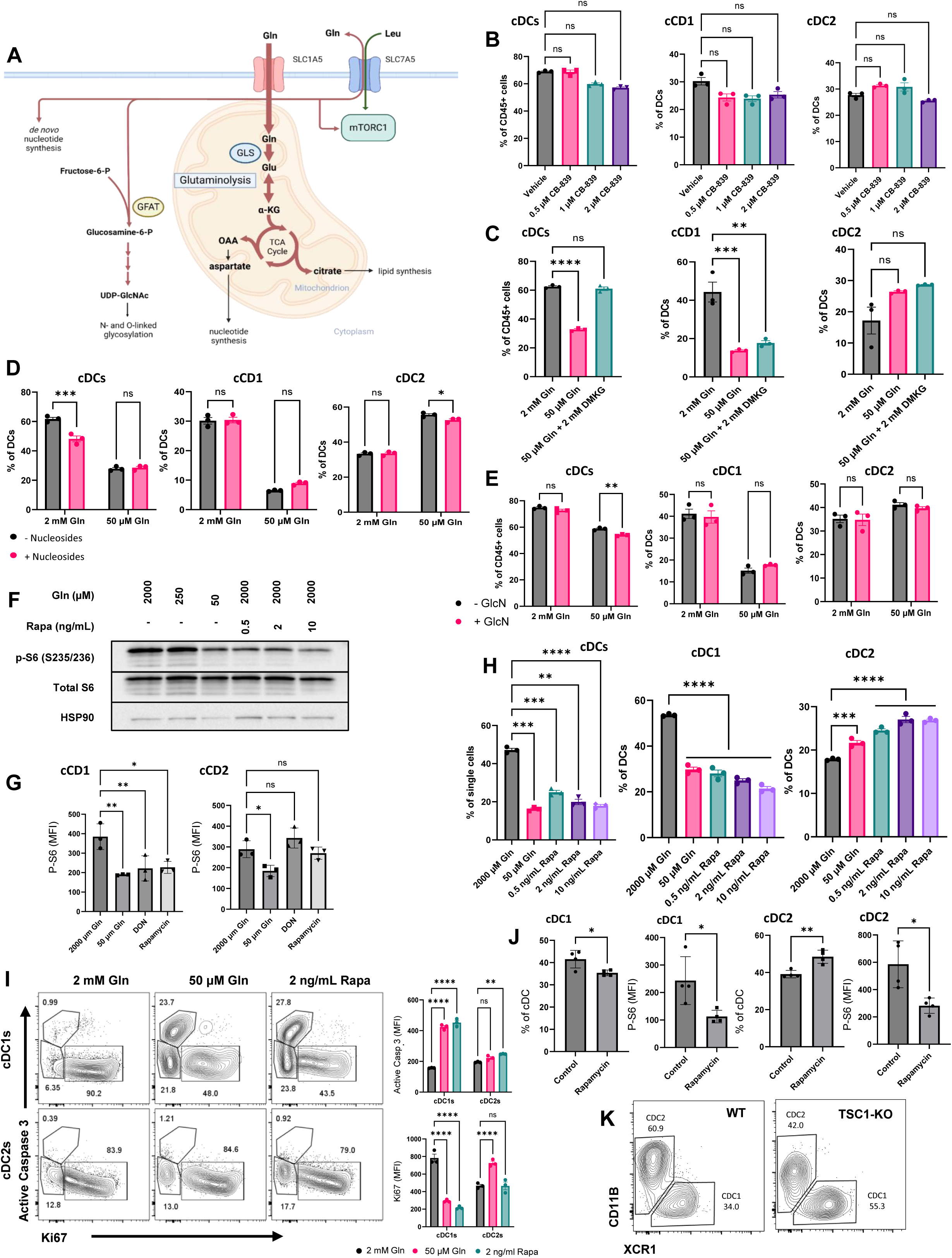
Glutamine regulates cDC1 frequency through MTOR signaling A: Basic overview of intracellular glutamine metabolism. B: Bar graphs quantifying the proportion of FLT3L-derived BMDCs (leftmost panel), defined as CD45^+^ B220^-^ *Zbtb46-GFP*^+^ cells, among CD45^+^ cells, and cDC1s and cDC2s (middle and right panels), defined above, among cDCs via flow cytometry after culture with the specified doses of the GLS antagonist CB-839 for 9 days. C: Bar graphs quantifying the proportion of FLT3L-derived BMDCs (leftmost panel), as defined in Figure 6B, among CD45^+^ cells, and cDC1s and cDC2s (middle and right panels) among cDCs via flow cytometry after culture with either 2 mM Gln, 50 μM Gln, or 50 μM Gln with 2 mM dimethyl 2 oxo-glutarate for 9 days. D: Bar graphs quantifying the proportion of FLT3L-derived BMDCs (leftmost panel), as defined in Figure 6B, among CD45^+^ cells, and cDC1s and cDC2s (middle and right panels) among cDCs via flow cytometry after culture with either 2 mM Gln or 50 μM Gln +/- 50 μM each of adenosine, cytidine, guanosine, thymidine, and uridine for 9 days. E: Bar graphs quantifying the proportion of FLT3L-derived BMDCs (leftmost panel), as defined in Figure 6B, among CD45^+^ cells, and cDC1s and cDC2s (middle and right panels) among cDCs via flow cytometry after culture with either 2 mM Gln or 50 μM Gln +/- 50 μM of glucosamine for 9 days. F: Western blot assessing phosphorylation status of S6 ribosomal protein compared to total S6 and HSP90 levels in FLT3L-driven BMDCs after 5 days of culture with 2 mM Gln, 250 μM Gln 50 μM Gln, or 0.5, 2, or 10 ng/mL of rapamycin. G: Phosphorylated S6 ribosomal protein measured via flow cytometry within cDC1 and cDC2 fraction of FLT3L-derived BMDCs after 7 days of culture with 2 mM Gln, 50 μM Gln, DON, or 10ng/ml Rapamycin. Y axis show median fluorescent intensity H: Bar graphs quantifying the proportion of FLT3L-derived BMDCs (leftmost panel), defined as CD45^+^ B220^-^ *Zbtb46-GFP*^+^ cells, among CD45^+^ cells, and cDC1s and cDC2s (middle and right panels), defined above, among cDCs via flow cytometry after culture for 7 days with the indicated concentrations of glutamine or doses of rapamycin. I: Representative flow cytometric plots and bar graphs quantifying the expression levels of Ki67 and active Caspase 3 among cDC1s, defined as CD45^+^ B220^-^ MHCII^+^ CD24^+^ CD11b^-^ cells, and cDC2s, defined as CD45^+^ B220^-^ MHCII^+^ CD24^+^ CD11b^-^ cells, after culture for 7 days with either 2 mM Gln, 50 μM Gln, or 2 mM Gln with 2 ng/mL rapamycin. J: Mice were treated with 8 mg/kg/day Rapamycin for 7 days followed by flow cytometric analyses of Lung DCs. cDCs were first identified as CD45^+^ CD11C^+^ CD64^-^ MHCII^+^ cells, within which cDC1 were CD103^+^ CD11b^-^ while cDC2 were CD103 CD11b^+^. K: Conditional *Tsc1^Flox/Flox^* mice were bred to *Rosa26^CAG-tdTomato^* reporter mice and the *Ms4a3^Cre^* mice to generate *Tsc1^Flox/Flox^ Rosa26^CAG-tdTomato^ Ms4a3^Cre^* (Tsc1-KO) and *Rosa26^CAG-tdTomato^ Ms4a3^Cre^* (WT). Shown are representative flow cytometry plots of lung DCs (CD45^+^ CD11C^+^ CD64^-^ MHCII^+^ cells, see Supplementary Figure 6A for gating).

Outside of its role in glutaminolysis, glutamine also contributes to other critical cellular pathways that sustain cellular growth, proliferation, and signaling. Glutamine is a critical nitrogen donor for *de novo* purine and pyrimidine synthesis (39). Additionally, in combination with products of glycolysis, glutamine is required to produce substrates for protein and lipid glycosylation via the hexosamine biosynthetic pathway (**Figure 6A**) (40). To investigate whether flux through these two pathways regulates cDC1 production, we separately added nucleosides (**Figure 6D**) and glucosamine (**Figure 6E**) to glutamine-deprived FLT3L-BMDC cultures. However, neither of these metabolites was able to rescue either total cDC or cDC1s in glutamine-deprived cultures.

An orthogonal study by Guo and colleagues (while our work was underway) showed that low glutamine levels in the tumor microenvironment impair cDC1 function without affecting cDC1 frequency (41). Here, the authors engendered a milder level of glutamine deficiency and shorter duration of cDC exposure to low glutamine availability. Indeed, we find a direct correlation between cDC1 frequency and glutamine levels where less dramatic reduction in glutamine has a limited impact on the frequency and numbers of cDC subsets (**Figures 5A-B**). Thus, falling glutamine levels likely impact cDC1 functions first and then reduce their numbers with increasing magnitude and duration of glutamine deficiency. Guo et al. found a role for FLCN and TFEB in mediating the functional impact of low glutamine on cDC1s. As our focus was on DC numbers, as opposed to function, we explored other potential mechanisms. Having ruled out multiple metabolic pathways above, we next explored the mTORC1 signaling complex as previous studies support its role in regulating cDC1 differentiation and survival (42, 43). Indeed, glutamine is known to be critical for mTORC1 activation via Rag-dependent and Rag-independent pathways (42, 44, 45). However, mTORC1 is activated by several essential and nonessential amino acids (for example leucine and arginine) (46) . In order to determine whether glutamine deprivation alone inhibits mTORC1 signaling, BMDCs were cultured at low glutamine levels or in the presence of rapamycin for several days and then phosphorylation of the downstream mTORC1 target S6 ribosomal protein was assessed (**Figure 6F**). Notably, cDC1s showed a greater reduction of Phospo-S6 under low glutamine or rapamycin treatment compared to cDC2s (**Figure 6G**). Correspondingly, rapamycin treatment selectively reduced cDC1 frequency in FLT3-BMDC cultures and was associated with reduced proliferation and increased apoptosis in cDC1s similar to low glutamine conditions (**Figures 6H-I**). Importantly, *in vivo* treatment with rapamycin reduced while conditional deletion of Tsc1 (inhibits mTORC) increased cDC1s in lungs (**Figure 6J-K**). These findings are consistent with a previous report demonstrating the requirement of mTORC1 for cDC1 specification in this tissue (42). Taken together, these results demonstrate that glutamine levels differentially regulate cDC subsets via modulation of mTORC1 signaling, at least in part.

## DISCUSSION

Inhibition of glutamine metabolism, especially using DON prodrugs in combination with immune checkpoint blockade, shows good promise for the treatment of diverse cancer types (13, 47, 48). Our work provides new insights into how these therapies may interact with DCs – a key regulator of adaptive immune responses.

Notably, a recent publication by Guo and colleagues asked similar questions regarding the role of glutamine in the tumor microenvironment and found cDC1 functions to be impaired when glutamine levels drop in this context (41). Our findings expand the scope of glutamine’s impact on the DC lineage beyond Guo et. al’s report and the two studies in parallel firmly establish a pivotal role for glutamine in this key antigen presenting cell.

Previously, Sinclair et al. generated a CD11c-specific *mTOR* deletion and observed that, similar to our results, lung cDC1s were significantly reduced (42). Our efforts link glutamine availability to this dependence on mTOR for cDC1 differentiation, proliferation, and survival in both normal tissue and tumors. This paints an emerging picture where FLCN/TFEB mediated functional defects within cDC1s (as described by Guo et. al.,) precede an mTOR-mediated drop in cDC1 frequency (described in this study) as glutamine levels continue to drop and its deficiency becomes more persistent. This ties nicely with the well-known dysfunction and paucity of cDC1s in tumors, which is one reason why solid tumors evade immune responses. In the context of immunotherapy, however, Leone and colleagues reported enhanced anti-tumor activity of T cells in response to DON (13). This ‘therapeutic conundrum’ regarding the use of DON as a drug might be resolved by targeting DON to specific cellular compartment or staggering the timing of DON treatment with other therapeutic modulation. An an example, DON could be targeted specifically to tumor cells through new formulations and restricted temporally (few days or weeks). As intratumoral CDCs have a limited life span (∼ one week) requiring continuous recruitment of precursor cells, cessation of DON treatment will then enable ‘re-population’ of the TME by cDC1s that can sample neo-antigens from tumor cells killed by DON. T cell responses to these neo-antigens can be further bolstered by subsequent treatment with immune checkpoint blockade.

The metabolic profile of solid tumors is an important consideration for the use of glutamine modulation as a therapeutic approach. Guo and colleagues employed colorectal carcinoma tumor cells (MC38) where intratumoral glutamine supplementation led to enhanced cDC1 numbers and increased anti-tumor T cell activity. In contrast, glutamine supplementation increased tumor growth in our sarcoma models (not shown). This is very likely due to the greater avidity of sarcomas for glutamine (6) where its supplementation enhances tumor cell proliferation and growth, thereby negative any anti-tumor immune effects. Thus, the optimal approach for therapeutic modulation of glutamine metabolism for cancer therapy is predicated on better understanding of how glutamine regulates tumor cells themselves *vs*. various stromal components including immune cells.

## MATERIALS AND METHODS

### Animals

C57BL/6 mice (4-6 weeks old, females) were purchased from Jackson Laboratories. *LSL-Kras^G12D/+^;Trp53^fl/fl^* mice were a generous gift from Dr. Tyler Jacks (31). *Zbtb46^GFP^*mice were a generous gift from Dr. Kenneth Murphy (32). Tsc1^fl/fl^ , Rosa26^CAG-tdTomato^ , and Ms4a3^Cre^ mice were purchased from the Jackson laboratories. Mice were bred and maintained in specific pathogen free facilities at the University of Pennsylvania. All animal procedures were conducted according to National Institutes of Health guidelines and approved by the Institutional Animal Care and Use Committee at the University of Pennsylvania.

### Cell lines

C57BL/6 syngeneic fibrosarcoma cell line was a generous gift from Dr. Robert Schrieber and was used for experimentation as previously described (49). Syngeneic UPS cell line was a generous gift from Dr. Sandra Ryeom. Fibrosarcoma-OVA-ZSGreen cell line was previously generated via lentiviral infection by Samir Devalaraja. Tumor cell lines were cultured in DMEM with 10% FBS, 1% Pen/Strep and 2mM glutamine. Low passage (< P20) cell lines were used for *in vitro* and *in vivo* experimentation. All cells were confirmed to be negative for mycoplasma contamination as assessed by MycoAlert Mycoplasma Detection Kit (Lonza).

### Tumor models

As previously described, tumors were generated in 8 weeks or older male and female LSL-Kras^G12D/+^;Trp53^fl/fl^ mice by injection of 30 units of Cre protein (TAT-Cre, Millipore) into the hindlimb musculature to minimize immune response to virus. Syngeneic fibrosarcoma flank tumors were established in C57BL/6 mice. shScramble or shGLS fibrosarcoma tumors were established in C57BL/6 or Zbtb46^GFP^ mice. As previously described, cultured tumor cells were detached using 0.05% trypsin (GIBCO), washed once with DMEM media and once with 1x PBS, and counted in preparation for implantation. Tumor cells were propagated *in vitro* for two passages prior to implantation and injected cells were greater than 90% viable. 5x10^5^ tumor cells were implanted subcutaneously (s.c.) into shaved flanks of recipient mice. Tumor dimensions were measured using a caliper starting at Day 6 and every two-three days thereafter. Animals were euthanized by CO2 inhalation at endpoint. Transplantable tumor volumes were calculated using the formula volume = (length × (width)^2^)/2. Sex (both male and female) and aged matched mice between 6-12 weeks of age were used for these studies.

### shRNA-mediated gene knockdown

pLKO.1 plasmids containing shRNAs targeting GLS or scramble control were purchased from Millipore-Sigma (SHCLND). As previously described, the vector was transfected into 293T cells using polyethylenimine (PEI) along with lentivirus packaging plasmids. Lentivirus supernatant was collected 48 hours later and passed through a 40uM filter. Subsequently, tumor cells were transduced and selected on 2ug/mL of puromycin for one week. Knockdown efficiency was determined by RT-qPCR for GLS, as well as assessment of *in vitro* glutamine uptake using a YSI metabolite analyzer.

### Measurement of glutamine uptake and glutamate production

Cells in culture were scraped and resuspended in DMEM (10% FBS, 1% P/S, 1% Gln). Cells were plated at a density of 200,000 cells/well in 6-well plates in a total of 3 mL of culture media. Wells with no cells were kept as controls for normalization of metabolite concentrations in media. After 24 hours post-seeding, media was removed, wells were washed with 1X DPBS, and fresh media was added to each well. Samples of culture media were collected 24 h post-media change and stored at −80°C until analysis after a quick spin down to eliminate contamination by cells and cell debris. Quantification of glutamine and glutamate concentrations (nM) in samples of cell culture media supernatant was determined enzymatically with a bioanalyzer (YSI2950, YSI Incorporated). For normalization purposes, number of cells per well was determined using a Countess III. Rate of metabolite consumption was calculated as previously published (50).

### In vivo treatment

As previously described, in tumor-bearing animals, once transplantable tumors were established (∼100 mm^3^) or hindlimb differences were observed (∼7 weeks post-injection), animals were randomized into two groups and treated daily with vehicle or drug delivered intraperitoneally. DON (Cayman Chemical Company 17580) was diluted in PBS and delivered daily at 0.25 mg/kg or 0.5mg/kg. V9302 (SelleckChem # S8818) was diluted in 10% DMSO, 45% PBS, and 45% PEG400 and delivered daily at 25 mg/kg. 200 μg of anti-CD3 monoclonal blocking antibody (clone 17A2, BioXCell) was administered intraperitoneally every three days starting on the day of implantation of shScramble or shGLS syngeneic fibrosarcoma cells. 10 μg of recombinant human/human FLT3L-Ig (BE0342, BioXCell) was administered intraperitoneally daily for duration of treatment.

### Tissue harvest and processing

As previously described, tissues were harvested and minced at indicated time points for analysis. Single cell suspensions were generated by digestion with collagenase B and DNase I for 45-60 minutes at 37° C and filtration through 70 µM cell strainer. Mouse blood was collected in EDTA tubes and RBCs were lysed using ACK lysing buffer, then quantified on a Countess II.

### Flow cytometry

Extracellular staining: Samples were harvested, resuspended in 100 μl of MACS buffer (PBS with 0.5% BSA and 2 mM EDTA), then incubated for 5 minutes on ice with anti-mouse CD16/32 Fc Block (BD, Clone 2.4G2, # 553142) at 1:200, and subsequently stained at 4 C with primary-fluorophore conjugated antibodies listed below for 20 minutes at 4 C for identification of cell populations by FACS. Intracellular staining for p-S6, Ki67, and Active Caspase 3 was performed using the Invitrogen FOXP3/Transcription factor staining kit (Cat # 00-5523- 00). Briefly, after extracellular staining, cells were incubated in Fix/Permeabilization buffer for 60 minutes at room temperature, then washed twice, incubated with Permeabilization buffer for 15 minutes, then stained with intracellular antibodies at 4 C overnight up to 16 hours, before being washed twice in Permeabilization buffer, then once in MACS buffer, then resuspended in MACS buffer. Flow cytometry was performed on an LSR Fortessa Flow Cytometer (BD Biosciences) and analyzed using FlowJo software (Treestar).

### Flow Cytometric Antibodies

All antibodies were used at 1:200 unless otherwise noted.

Biolegend: CD45R/B220 – APC/Cy7 (Clone RA3-6B2, # 103224), BV510 (Clone RA3-6B2, # 103247); CD8α – AF647 (Clone 53-6.7, # 100724), PE/Dazzle 594 (Clone 53-6.7, # 100762); CD11c – APC/Cy7 (Clone N418, # 11), PE-Dazzle-594 (Clone N418, # 117348); CD24 – APC (Clone M1/69, # 101814); CD45 – BV510(Clone IM7, # 103043), BV605 (Clone 30-F11, # 103139); CD64 – PE (Clone X54-5/7.1, #139304); CD103 – AF647 (Clone 2E7, #121410), PE/Dazzle 594 (Clone 2E7, #121430); CD172α/SIRPα – PerCP/Cy5.5 (Clone P84, # 144009), BV510 (Clone P84, #144302), FITC (Clone P84, # 144006); I-A/I-E 1:400 – BV421 (Clone M5/114.15.2, # 107631); Ly6c – BV605 (Clone HK1.4, #128036), PE (Clone HK1.4, #128007); Ly6g 1:400 – PerCP/Cy5.5 (Clone 1A8, # 127616); F4/80 – BV605 (Clone BM8, # 123133); XCR1 – BV510 (Clone ZET, # 148218) Invitrogen: CD11b 1:400 – PE/Cy7 (Clone M1/70, # 25-0112-82), BUV805 (Clone M1/70, # 368-0112-82) BD: CD45 - BUV395 (Clone 30-F11, # 564279); CD135 1:100 for 60 minutes at 4 C – PE-CF594 (Clone 562537, # A2F10.1); CD117 1:100 for 60 minutes at 4 C – BUV395 (Clone 2B8, # 564011); Ki67 stained intracellularly – PE/Cy7 (Clone B56, # 561283); Active Caspase 3 stained intracellularly – FITC (Clone C92-605, # 560901) Cell signaling: P-S6 – PE (Clone D57.2.2E, # 5316S)

### Bone marrow isolation and culture

Femurs and tibias from indicated mice were dissected from mouse hindlimbs and flushed through with MACS buffer onto 70 micron strainer to remove marrow. RBCs were then lysed using ACK lysis buffer and remaining cells were quantified using a Countess II.

### FLT3L Culture

Isolated bulk bone marrow cells were cultured for 7-10 days in glutamine-free IMDM (Sigma-Aldrich I3390- 500ML) supplemented with 10% dialyzed FBS (Gemini Bio-Products 100-108), 1% Pen/Strep, 1% Sodium Pyruvate, 0.1% Beta-mercaptoethanol, 150 ng/mL FLT3L (Peprotech # 250-31L-250UG) and the indicated concentration of glutamine, as well as the indicated levels of drugs or metabolites. V9302, CB-839, and rapamycin were diluted in DMSO. For any experiments with metabolite addbacks, media was refreshed on days 3 and 6 of culture. Metabolites/pharmaceutical compounds used: Dimethyl 2-oxo-glutarate (Sigma-Aldritch 349631-5G), Glucosamine (Sigma-Aldrich G1514-100G), Adenosine (Sigma-Aldrich A4036-5G), Cytidine (Sigma-Aldrich C4654-1G), Guanosine (Sigma-Aldrich G6264-1G), Thymidine (Sigma-Aldrich T1895-1G), Uridine (Sigma-Aldritch U3003-5G), CB-839 (SelleckChem # S7655), Rapamycin (Stem Cell # 73362)

### Single Cell RNA Sequencing

#### Sample preparation, scRNA-sequencing and pre-processing

Control or DON-treated tumors (n = 1 per group) were harvested after 6 days of treatment and positively selected for CD45+ cells using Mouse CD45 Microbeads (Miltenyi Biotec) before submission. FLT3L-BMDCs were submitted for scRNA-seq without prior enrichment. Preparation of sequencing libraries and sequencing were conducted as previously described at the Center for Applied Genomics Core at the Children’s Hospital of Philadelphia (51). Pooled libraries were prepared using the 10x Genomics Chromium Single Cell 3′ Reagent kit v3 per manufacturer’s instructions. Sequencing was performed on either an Illumina HiSeq 2500 or NovaSeq 6000. Demultiplexing and alignment of sequencing reads to the mm10 transcriptome and creation of feature- barcode matrices was done using the CellRanger pipeline (10X Genomics, v.3.0.0/3.0.2/3.1.0/6.0.0).

#### Standard scRNA-seq analysis

As previously described, For each sample, the processed output files from CellRanger (barcodes,fgenes, matrix) were passed to the Seurat package (v.4.3.0) for downstream processing in R (v.4.3.1). Genes expressed in less than 3 cells were filtered out. Outlier cells were identified based on library size, number of expressed genes and mitochondrial proportion using the standard Seurat pipeline for quality control and were subsequently removed. Seurat pipeline was used to normalize data (NormalizeData) and identify highly variable genes (FindVariableFeatures) and principal component (PC) analysis was performed on the scaled 2000 most variable genes (RunPCA). The first 20 PCs were used for graph-based cluster identification (FindNeighbors, FindClusters) and UMAP dimensional reduction (RunUMAP). Clustering granularity was set at 0.5 resolution for clusters shown in Figures X, Y, Z. Differentially expressed genes (DEGs) were identified using the default Wilcoxon rank sum test in the FindAllMarkers and FindMarkers functions from Seurat.

#### Annotation of cell identities by SingleR

Annotation of cell identities based on reference data was performed by SingleR (v.2.2.0). To annotate the UPS CD45+ and FLT3L-BMDC datasets (Fig. X, Y, Z), cell identities were assigned using ImmGen reference data. Normalized expression data of mouse immune cell types from the Immunological Genome Project (ImmGen) were downloaded via celldex (v.1.10.1).

#### RNA isolation and qPCR analysis for gene expression

Total RNA was isolated using GenElute Mammalian Total RNA Miniprep Kit (Sigma). Reverse transcription was performed using the High Capacity RNA to cDNA Kit (Life Technologies). qRT-PCR was performed using ViiA7 Real-Time PCR machine and TaqMan probes used for gene specific amplification (purchased from ThermoFisher Scientific). Probes: Hprt (Mm03024075_m1), Gls (Mm01257297_m1)

### Immunoblotting

As previously described, cells were harvested in lysis buffer (40 mM HEPES (pH 7.4), 2 mM EDTA, 10 mM pyrophosphate, 10 mM glycerophosphate, 1% Triton X-100) with Complete Ultra protease/phosphatase inhibitor (Roche; Cat. #05892791001). Samples were centrifuged at 20,000×g for 15 min at 4 °C. Protein lysates were resolved by Tris-Glycine SDS-PAGE and transferred to nitrocellulose membranes (Biorad; Cat. #162-0115). Membranes were incubated with primary antibodies overnight at 4 °C diluted in milk or TBST supplemented with 5% bovine serum albumin (BSA). Antibodies used for western blot: GLS 1:1000 (Abcam ab93434), GAPDH 1:1000 (Cell signaling # 5174), P-S6 1:2000 (Cell Signaling # 2211S), Total S6 1:1000 (Cell Signaling # 2217S), HSP90 1:1000 (Cell Signaling # 4874)

### Statistical Analysis

Statistical analyses were performed using GraphPad Prism 9 (GraphPad). T test was used for comparison of two groups and one-way ANOVA with Tukey’s HSD post-test was used for multiple comparisons. *p* < 0.05 was considered statistically significant (\**p* < 0.05, \*\**p* < 0.01, \*\*\**p* < 0.001, \*\*\*\**p* < 0.001) and error bars represent SD.

## Data Availability

All sequencing data described in this manuscript are being uploaded to GEO and made publicly available upon publication.

## Acknowledgements

We would like to acknowledge Dr. Brian Keith for his helpful comments and input. We would also like to thank the core facilities at the University of Pennsylvania and the Children’s Hospital of Philadelphia, especially the flow cytometry core at Upenn and the CAG sequencing core at CHOP. We would also like to acknowledge the funding sources that includes R01 DK138827 (M.H. and M.C.S), CRI STAR award (M.H), F30 CA265069 (G.L), and F31 CA261041 (M.C.N).

## Author Contributions

Conception and design: G.L., N.H., M.C.S., and M.H.

<u>Development of methodology</u>: G.L., N.H., W.A.MA., M.S., M.C.J., M.C.S., and M.H.

<u>Acquisition of data</u>: G.L., N.H., W.A.MA., M.S., J.S., M.C.N., J.T., L.Z., N.P.L., and M.H

<u>Analysis and interpretation of data</u>: G.L., N.H., W.A.MA., M.S., J.S., M.C.N., M.C.S., and M.H.

Writing, review, and/or revision of the manuscript: G.L., N.H., W.A.MA., J.S., M.C.S., and MH

Study supervision: M.C.S., and M.H.

## Competing interests

The authors declare no competing interest.

| Reagent & Resource | Source | Identifier |
| --- | --- | --- |
| **Antibodies** |  |  |
| Anti-Glutaminase antibody | Abcam | Cat#ab93434 |
| Anti-Active Caspase-3 (C92-605) FITC | BD Biosciences | Cat#560901 |
| Anti-mouse CD117 (2B8) BUV395 | BD Biosciences | Cat#564011 |
| Anti-mouse CD135 (562537) PE-CF594 | BD Biosciences | Cat#A2F10.1 |
| Anti-mouse CD16/32 Fc Block | BD Biosciences | Cat#553142 |
| Anti-mouse CD45 (30-F11) BUV395 | BD Biosciences | Cat#564279 |
| Mouse Ki-67 (B56) PE/Cy7 | BD Biosciences | Cat#561283 |
| InVivoMAb anti-mouse CD3 | Bio X Cell | Cat#BE0002 |
| InVivoMAb recombinant Flt-3L-Ig (hum/hum) | Bio X Cell | Cat#BE0342 |
| Anti-mouse CD103 (2E7) AF647 | BioLegend | Cat#121410 |
| Anti-mouse CD103 (2E7) PE/Dazzle 594 | BioLegend | Cat#121430 |
| Anti-mouse CD11c (N418) APC/Cy7 | BioLegend | Cat#11 |
| Anti-mouse CD11c (N418) PE-Dazzle-594 | BioLegend | Cat#117348 |
| Anti-mouse CD172a/SIRPα (P84) BV510 | BioLegend | Cat#144302 |
| Anti-mouse CD172a/SIRPα (P84) FITC | BioLegend | Cat#144006 |
| Anti-mouse CD172a/SIRPα (P84) PerCP/Cy5.5 | BioLegend | Cat#144009 |
| Anti-mouse CD24 (M1/69) APC | BioLegend | Cat#101814 |
| Anti-mouse CD45 (30-F11) BV605 | BioLegend | Cat#103139 |
| Anti-mouse CD45 (IM7) BV510 | BioLegend | Cat#103043 |
| Anti-mouse CD45R/B220 (RA3-6B2) APC/Cy7 | BioLegend | Cat#103224 |
| Anti-mouse CD45R/B220 (RA3-6B2) BV510 | BioLegend | Cat#103247 |
| Anti-mouse CD64 (X54-5/7.1) PE | BioLegend | Cat#139304 |
| Anti-mouse CD8α (53-6.7) AF647 | BioLegend | Cat#100724 |
| Anti-mouse CD8α (53-6.7) PE/Dazzle 594 | BioLegend | Cat#100762 |
| Anti-mouse F4/80 (BM8) BV605 | BioLegend | Cat#123133 |
| Anti-mouse I-A/I-E (M5/114.15.2) BV421 | BioLegend | Cat#107631 |
| Anti-mouse Ly6c (HK1.4) BV605 | BioLegend | Cat#128036 |
| Anti-mouse Ly6c (HK1.4) PE | BioLegend | Cat#128007 |
| Anti-mouse Ly6g (1A8) PerCP/Cy5.5 | BioLegend | Cat#127616 |
| Anti-mouse/rat XCR1 (ZET) BV510 | BioLegend | Cat#148218 |
| GAPDH (D16H11) XP® Rabbit mAb | Cell Signaling | Cat#5174 |
| HSP90 Antibody | Cell Signaling | Cat#4874 |
| Phospho-S6 Ribosomal Protein (Ser235/236) (D57.2.2E) PE | Cell Signaling | Cat#5316S |
| Phospho-S6 Ribosomal Protein (Ser235/236) Antibody | Cell Signaling | Cat#2211 |
| S6 Ribosomal Protein (5G10) Rabbit mAb | Cell Signaling | Cat#2217S |
| Anti-mouse CD11b (M1/70) BUV805 | Invitrogen | Cat#368-0112-82 |
| Anti-mouse CD11b (M1/70) PE/Cy7 | Invitrogen | Cat#25-0112-82 |
| **Cell Culture Reagents** |  |  |
| Dialyzed Fetal Bovine Serum | Gemini Bio | Cat#100-108 |
| Recombinant Murine Flt3-Ligand | PeproTech | Cat#250-31L |
| Iscove's Modified Dulbecco's Medium | Sigma-Aldrich | Cat#I3390 |
| 2-Mercaptoethanol | Sigma-Aldrich | Cat#M3148 |
| DMEM, high glucose, GlutaMAX™ Supplement, pyruvate | Thermo Fisher Scientific | Cat#10569010 |
| DPBS, 1X | Thermo Fisher Scientific | Cat#MT21030CM |
| L-Glutamine | Thermo Fisher Scientific | Cat#25030081 |
| Penicillin-Streptomycin (10,000 U/mL) | Thermo Fisher Scientific | Cat#15140122 |
| Trypsin-EDTA (0.05%) | Thermo Fisher Scientific | Cat#25300054 |
| **Drugs** |  |  |
| Rapamycin | Cayman Chemical Company | Item no.13346 |
| 6-diazo-5-oxo-L-nor-Leucine (DON) | Cayman Chemical Company | Item no.17580 |
| V-9302 | SelleckChem | Cat#S8818 |
| Telaglenastat (CB-839) | SelleckChem | Cat#S7655 |
| Rapamycin | STEMCELL Technologies | Cat#73362 |
| **Experimental Model: Cell Lines** |  |  |
| UPS | Dr. Sandra Ryeom | N/A |
| C57BL/6 syngeneic fibrosarcoma | Dr. Robert Schrieber | N/A |
| Fibrosarcoma-OVA-ZSGreen | Dr. Samir Devalaraja | N/A |
| 293T |  |  |
| **Experimental Model: Strains** |  |  |
| *LSL-Kras<sup>G12D/+</sup>;Trp53<sup>fl/fl</sup>* | Dr. Tyler Jacks | N/A |
| C57BL/6 | Jackson Laboratories | N/A |
| *Ms4a3<sup>Cre</sup>* | Jackson Laboratories | N/A |
| *Rosa26<sup>CAG-tdTomato</sup>* | Jackson Laboratories | N/A |
| *Rosa26<sup>CAG-tdTomato</sup>Ms4a3<sup>Cre</sup>* (WT) | Jackson Laboratories | N/A |
| *Tsc1<sup>fl/fl</sup>* | Jackson Laboratories | N/A |
| *Zbtb46<sup>GFP</sup>* | Dr. Kenneth Murphy | N/A |
| **Plasmids** |  |  |
| pLKO.1 | Millipore-Sigma? Addgene? | Cat#185596? |
| **Metabolites/pharmaceutical compounds** |  |  |
| Dimethyl 2-oxo-glutarate | Sigma-Aldrich | Cat#349631 |
| Glucosamine | Sigma-Aldrich | Cat#G1514 |
| Adenosine | Sigma-Aldrich | Cat#A4036 |
| Cytidine | Sigma-Aldrich | Cat#C4654 |
| Guanosine | Sigma-Aldrich | Cat#G6264 |
| Thymidine | Sigma-Aldrich | Cat#T1895 |
| Uridine | Sigma-Aldrich | Cat#U3003 |
| **Tissue harvest reagents** |  |  |
| Collagenase, Type II | Thermo Fisher Scientific | Cat#17-101-015 |
| Deoxyribonuclease I | Sigma-Aldrich | Cat#D4527 |
| RBC Lysis Buffer | Santa Cruz Biotechnology | Cat#sc-296258 |
| **Others** |  |  |
| Foxp3 / Transcription Factor Staining Buffer Set | Thermo Fisher Scientific | Cat#00-5523-00 |
| TAT-CRE Recombinase | Millipore | Cat#SCR508 |
| Puromycin | InvivoGen | Cat#ant-pr-1 |
| PEG300 | MedChem Express | Cat#HY-Y0873 |
| PEG400 | MedChem Express | Cat#HY-Y0873A |
| Dimethyl sulfoxide | Sigma-Aldrich | Cat#D2650 |
| CD45 MicroBeads, mouse | Miltenyi Biotec | Cat#130-052-301 |
| HEPES | Thermo Fisher Scientific | Cat#15630080 |
| EDTA | Thermo Fisher Scientific | Cat#15575020 |
| Pyrophosphate |  |  |
| Glycerophosphate |  |  |
| Triton X-100 | Sigma-Aldrich | Cat#X100 |
| Complete Ultra protease/phosphatase inhibitor | Roche | Cat#05892791001 |
| Nitrocellulose Membrane | Bio-Rad | Cat#1620115 |
| polyethylenimine |  |  |

| RNA isolation and qPCR analysis reagents |  |  |
| --- | --- | --- |
| GenElute Mammalian Total RNA Miniprep Kit | Sigma-Aldrich | Cat#RTN70 |
| High-Capacity RNA-to-cDNA™ Kit | Thermo Fisher Scientific | Cat#4387406 |
| TaqMan Probes |  |  |
| Hprt (Mm03024075_m1) |  |  |
| Gls (Mm01257297_m1) |  |  |

---

**Figure S1.**
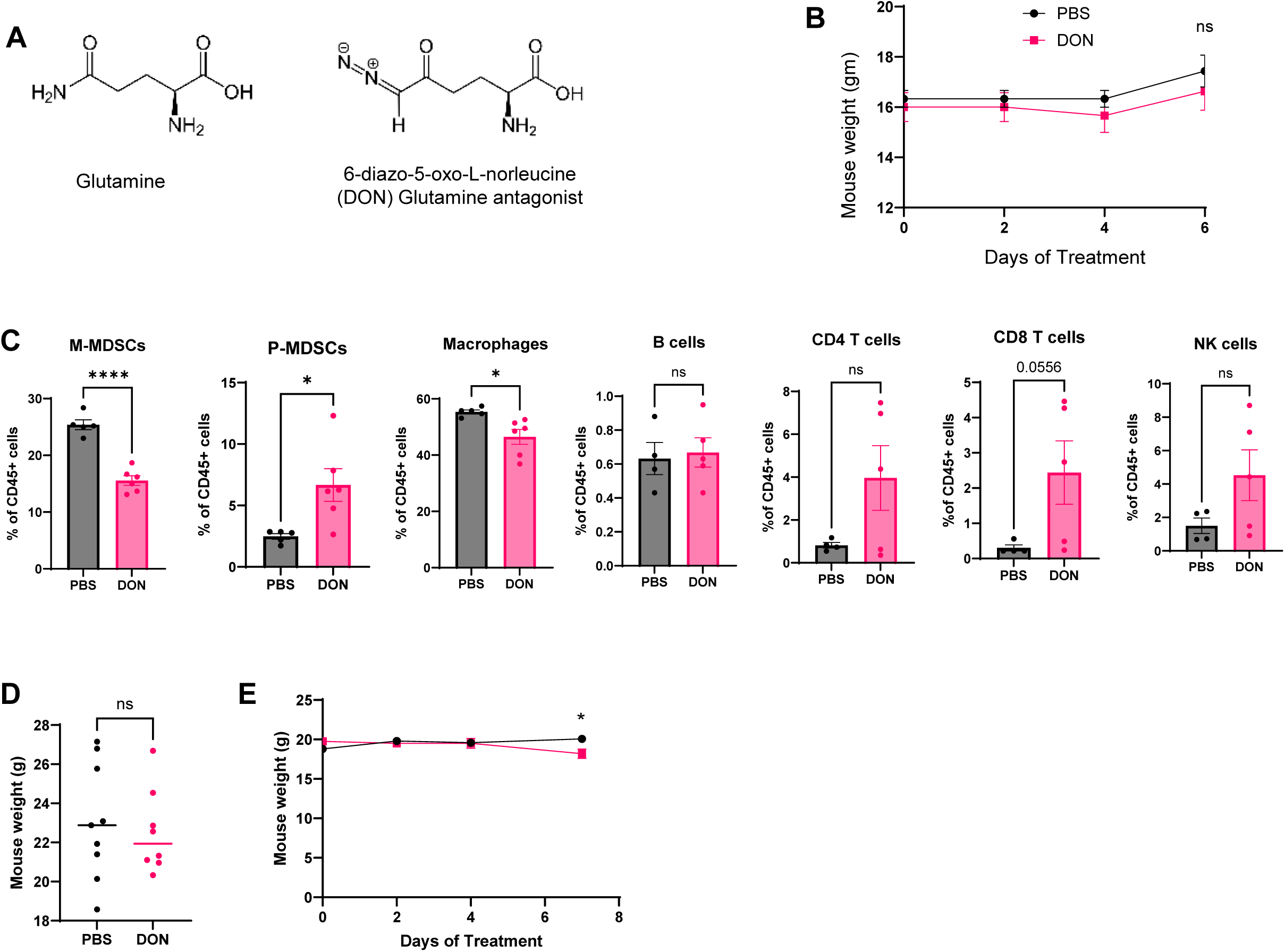
A: Structure of glutamine and the glutamine analog DON. B: Fibrosarcoma-bearing mouse weights over the course of PBS or DON treatment. C: Bar graphs quantifying the proportions of intratumoral M-MDSCs (defined as CD45+ CD11b+ Ly6g- Ly6c- hi cells), P-MDSCs (defined as CD45+ CD11b+ Ly6g+ Ly6c-lo cells), and macrophages (defined as CD45+ Ly6c- Ly6g- CD11b+ F4/80+ cells) among CD45+ cells in PBS or DON treated tumors. D: Weights of spontaneous LSL-Kras^G12D/+^;Trp53^fl/fl^ UPS tumor-bearing mice after 12 days of PBS or 0.25 mg/kg DON treatment. E: Weights of transplantable syngeneic UPS-bearing mice treated with either PBS or 0.5 mg/kg DON during the duration of treatment.

**Figure S2.**
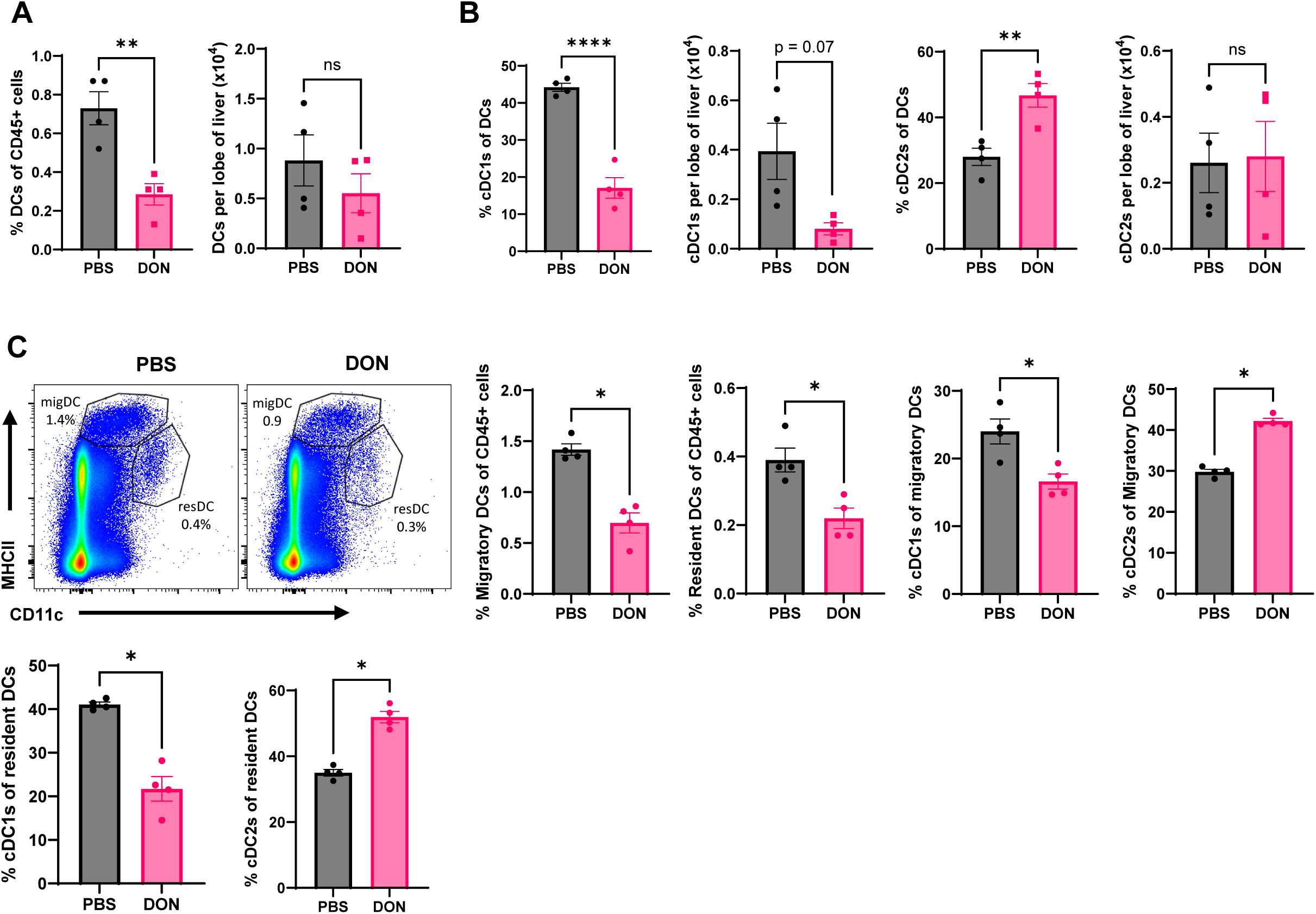
A: Bar graphs quantifying flow cytometric data of total liver DCs, defined as CD45+ Ly6c- Ly6g- CD64- CD11c+ MHCII+ cells, after PBS or DON treatment as a percentage of CD45+ cells (left graph) and per liver lobe (right graph). B: Bar graphs quantifying liver cDC1s and cDC2s, as defined in Figure 1C, after PBS or DON treatment as a percentage of DCs and per liver lobe. C: Representative flow cytometric plots quantifying resident (CD11C^high^MHCII^intermediate^) and migratory (CD11C^intermediate^MHCII^high^) DCs in the inguinal lymph nodes after treating mice for 10 days with PBS or DON. Bar graphs show migratory and resident DCs as a percentage of CD45+ cells, percentage of cDC1s (CD8α+ CD11b-) and cDC2 (CD8α- CD11b+) within resident DCs, and percentage of cDC1s (CD103+ CD11b-) and cDC2 (CD103- CD11b+) within migratory DCs under the two treatment conditions.

**Figure S3.**
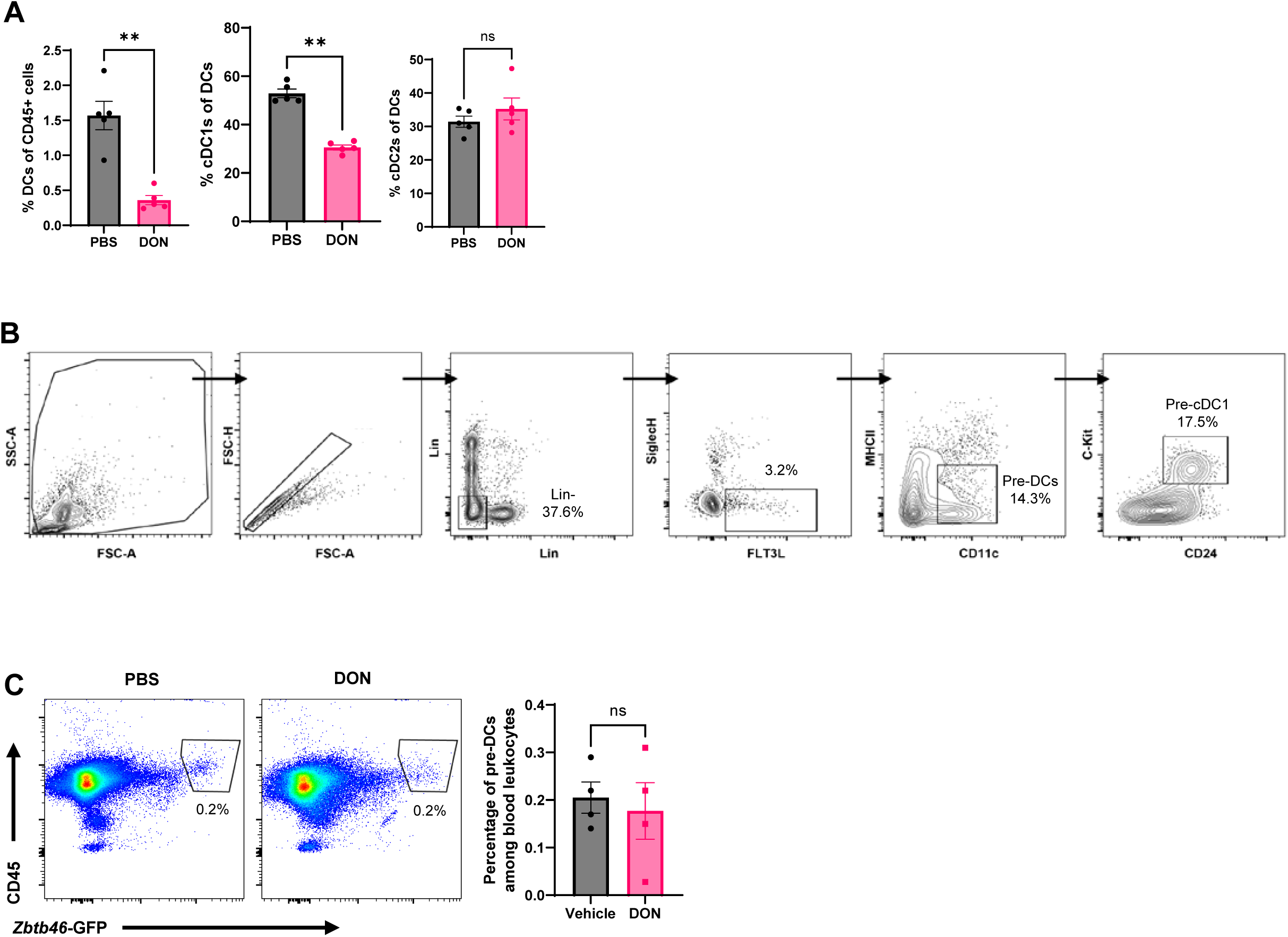
A: Bar graphs quantifying the percentage of lung DCs among all leukocytes (leftmost panel) and cDC1s and cDC2s among DCs (right two panels) after 6 days of daily DON (0.25 mg/kg) treatment. B: Gating scheme for the identification of bone marrow pre-cDCs and pre-cDC1s. C: Representative flow cytometric plots and bar graphs quantifying the proportion of CD45+ Zbtb46-GFP+ cells in blood as a percentage of CD45+ cells after 10 days of treatment with PBS or DON.

**Figure S4.**
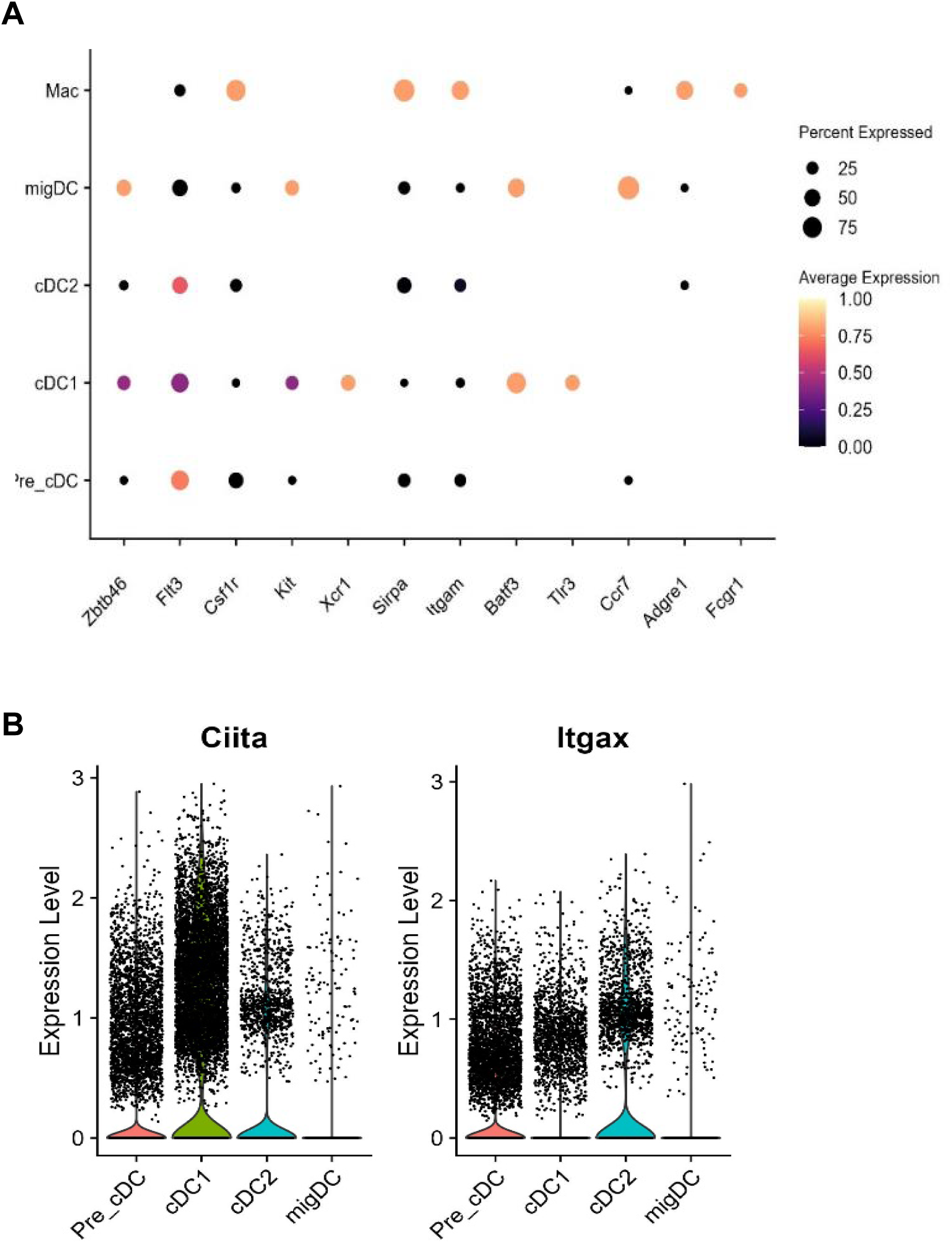
scRNA-seq (10X genomics) of FLT3 ligand induced bone-marrow derived DCs. (A) shows dot plots for the expression of key genes (X-axis) in the major DC subsets and macrophages (Y-Axis). (B) shows violin plots for the expression levels of the indicated genes in the major DC subsets.

**Figure S5.**
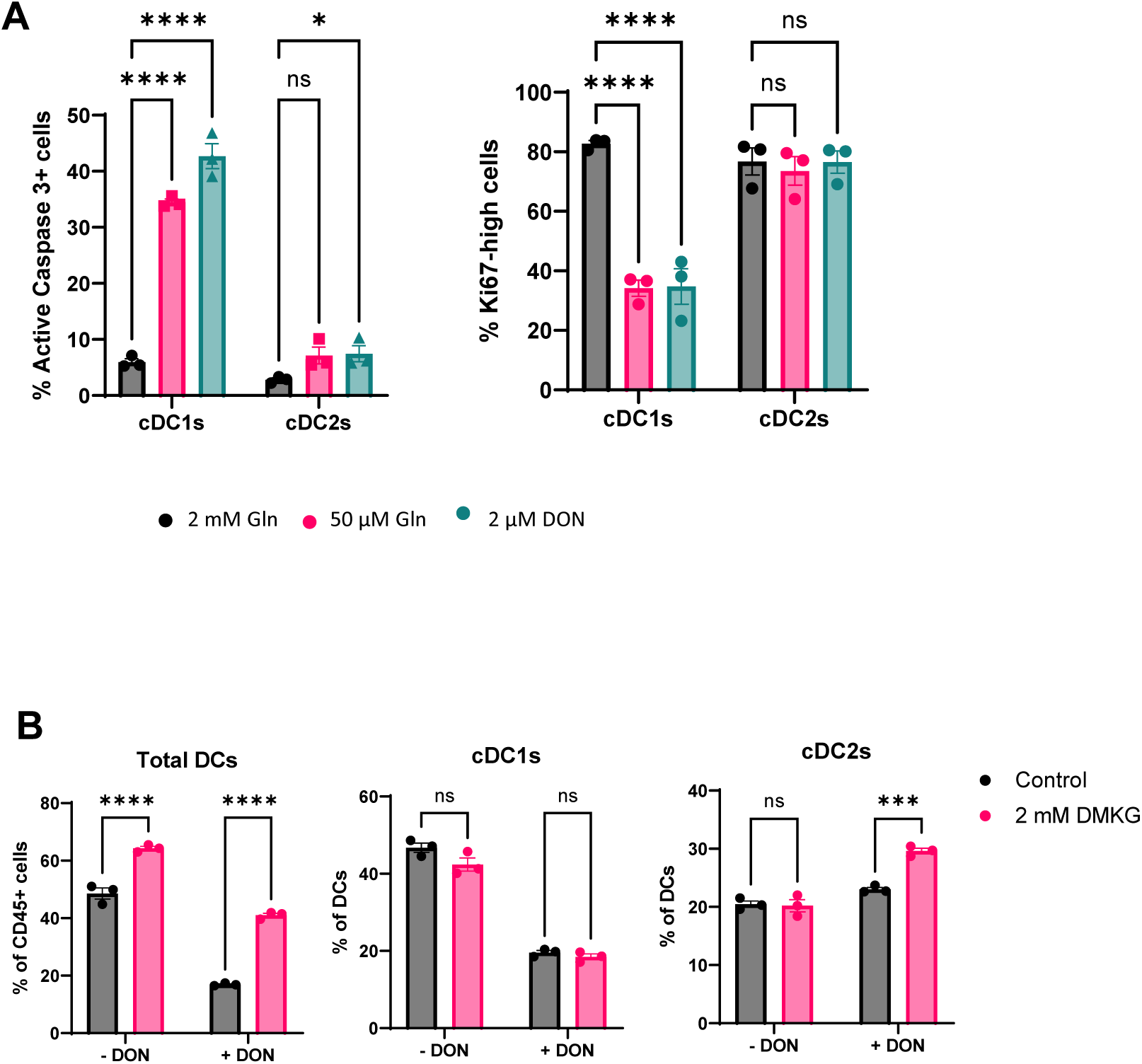
A: Bar graphs quantifying the proportions of Ki67+ Active Caspase 3- proliferating cells and Ki67- Active Caspase 3+ apoptotic cells among cDC1s and cDC2s after culture for 5 days with either 2 mM Gln, 50 μM Gln, or 2mM Gln with 2 μM DON. Please see Figure 5F for the corresponding representative flow cytometric plots. B: Bar graphs quantifying the proportion of FLT3L-derived BMDCs (leftmost panel), as defined in B), among CD45+ cells, and cDC1s and cDC2s (middle and right panels) among cDCs via flow cytometry after culture with either vehicle or or 1 μM DON +/- 2 mM dimethyl 2 oxo-glutarate for 9 days.

**Figure S6.**
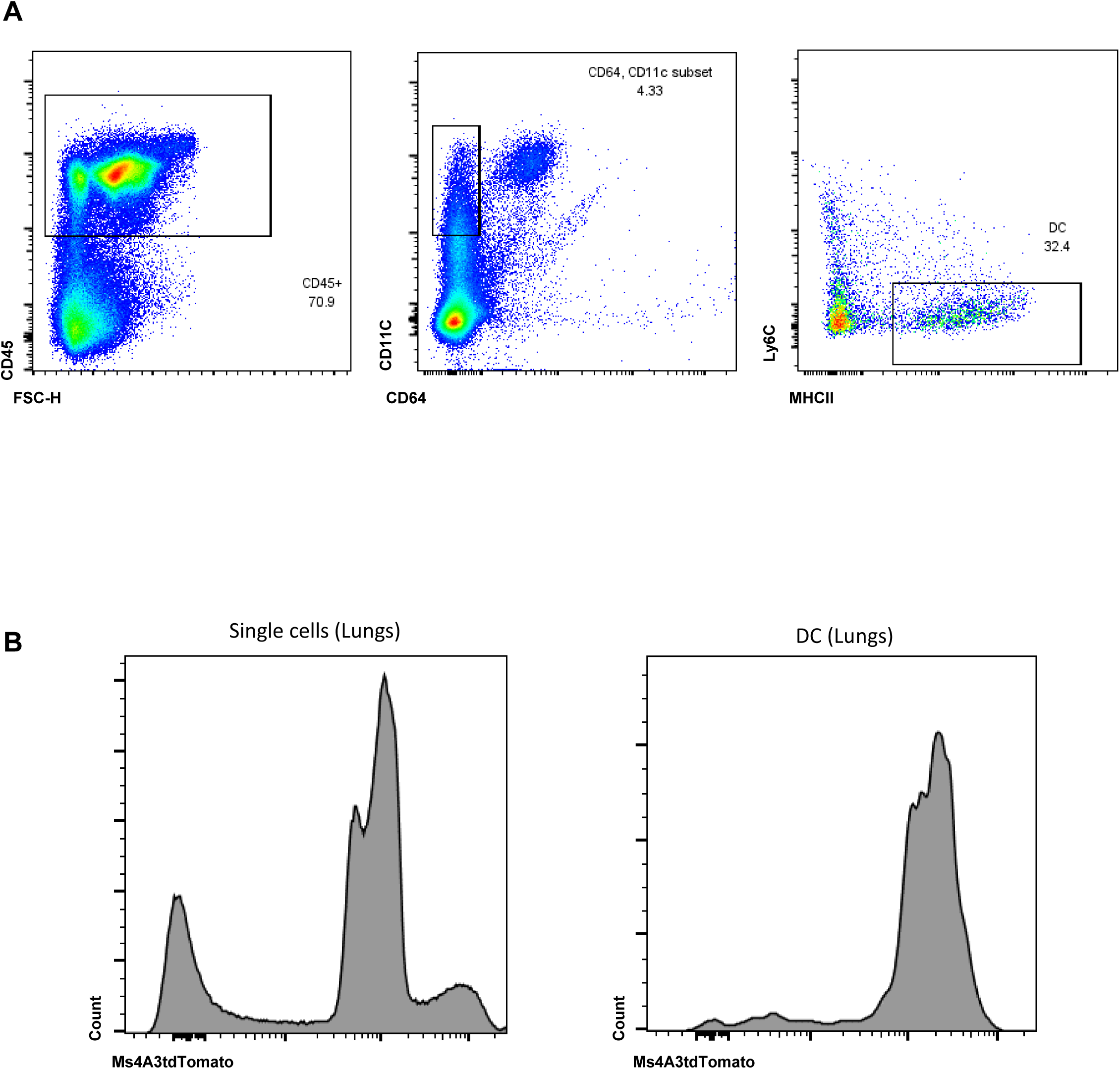
(A) Conditional Tsc1^Flox/Flox^ mice were bred to Rosa26^CAG-tdTomato^ reporter mice and the Ms4a3^Cre^ mice to generate Tsc1^Flox/Flox^ Rosa26^CAG-tdTomato^ Ms4a3^Cre^ (Tsc1-KO) and Rosa26^CAG-tdTomato^ Ms4a3^Cre^ (WT). Shown are representative flow cytometry plots displaying the gating strategy to identify lung DCs (see also Figure 6K). (B) Histogram showing expression of the Ms4A3 lineage via tdTomato expression in all lung cells (left) and specifically within DCs (right) identified using the gating scheme described above. The data shows all DCs being derived from Ms4A3 lineage and hence should lack Tsc1 when this mouse is bred with Tsc1-floxed mice.

## References

1. B. J. Altman, Z. E. Stine, C. V. Dang, From Krebs to clinic: glutamine metabolism to cancer therapy. Nat. Rev. Cancer 16, 619–634 (2016).

2. K. M. Lemberg, J. J. Vornov, R. Rais, B. S. Slusher, We’re Not “DON” Yet: Optimal Dosing and Prodrug Delivery of 6-Diazo-5-oxo-L-norleucine. Mol. Cancer Ther 17, 1824–1832 (2018).

3. M. I. Gross, Antitumor Activity of the Glutaminase Inhibitor CB-839 in Triple-Negative Breast Cancer. Mol Cancer Ther 13, 890–901 (2014).

4. R. M. Thompson, Glutaminase inhibitor CB-839 synergizes with carfilzomib in resistant multiple myeloma cells. Oncotarget 8, 35863–35876 (2017).

5. J. Zhang, Inhibition of GLS suppresses proliferation and promotes apoptosis in prostate cancer. Biosci Rep 39 (2019).

6. P. Lee, Targeting glutamine metabolism slows soft tissue sarcoma growth. Nature Communications 11, 1–15 (2020).

7. C.-H. Lee, Telaglenastat plus Everolimus in Advanced Renal Cell Carcinoma: A Randomized, Double-Blinded, Placebo-Controlled, Phase II ENTRATA Trial. Clin Cancer Res 28, 3248–3255 (2022).

8. N. M. Tannir, Efficacy and Safety of Telaglenastat Plus Cabozantinib vs Placebo Plus Cabozantinib in Patients With Advanced Renal Cell Carcinoma: The CANTATA Randomized Clinical Trial. JAMA Oncol 8, 1411–1418 (2022).

9. S. M. Davidson, Environment Impacts the Metabolic Dependencies of Ras-Driven Non-Small Cell Lung Cancer. Cell Metabolism 23, 517–528 (2016).

10. D. E. Biancur, Compensatory metabolic networks in pancreatic cancers upon perturbation of glutamine metabolism. Nature Communications 8, 1–15 (2017).

11. R. Rais, Discovery of 6-Diazo-5-oxo-l-norleucine (DON) Prodrugs with Enhanced CSF Delivery in Monkeys: A Potential Treatment for Glioblastoma. J. Med. Chem 59, 8621–8633 (2016).

12. M.-H. Oh, Targeting glutamine metabolism enhances tumor specific immunity by modulating suppressive myeloid cells. J Clin Invest (2020). 10.1172/JCI131859.

13. R. D. Leone, Glutamine blockade induces divergent metabolic programs to overcome tumor immune evasion. Science 366, 1013–1021 (2019).

14. M. Nakaya, Inflammatory T cell responses rely on amino acid transporter ASCT2 facilitation of glutamine uptake and mTORC1 kinase activation. Immunity 40, 692–705 (2014).

15. L. Araujo, P. Khim, H. Mkhikian, C.-L. Mortales, M. Demetriou, Glycolysis and glutaminolysis cooperatively control T cell function by limiting metabolite supply to N-glycosylation. eLife 6, 21330.

16. D. Klysz, Glutamine-dependent α-ketoglutarate production regulates the balance between T helper 1 cell and regulatory T cell generation. Sci Signal 8, **ra97** (2015).

17. Q. Yu, Targeting Glutamine Metabolism Ameliorates Autoimmune Hepatitis via Inhibiting T Cell Activation and Differentiation. Front Immunol 13, 880262 (2022).

18. M. O. Johnson, Distinct Regulation of Th17 and Th1 Cell Differentiation by Glutaminase-Dependent Metabolism. Cell 175, 1780–1795 19 (2018).

19. D. N. Edwards, Selective glutamine metabolism inhibition in tumor cells improves antitumor T lymphocyte activity in triple-negative breast cancer. J Clin Invest 131 (2021).

20. J.-K. Byun, Inhibition of Glutamine Utilization Synergizes with Immune Checkpoint Inhibitor to Promote Antitumor Immunity. Molecular Cell 80, 592–606 8 (2020).

21. W.-C. Wu, Immunosuppressive Immature Myeloid Cell Generation Is Controlled by Glutamine Metabolism in Human Cancer. Cancer Immunol Res 7, 1605–1618 (2019).

22. Q. Song, Dissecting intratumoral myeloid cell plasticity by single cell RNA-seq. Cancer Medicine 8, 3072–3085 (2019).

23. V. Durai, K. M. Murphy, Functions of Murine Dendritic Cells. Immunity 45, 719–736 (2016).

24. J. P. Böttcher, C. R., Sousa, The Role of Type 1 Conventional Dendritic Cells in Cancer Immunity. Trends in Cancer 4, 784–792 (2018).

25. D. Pakalniškytė, B. U. Schraml, “Chapter Three - Tissue-Specific Diversity and Functions of Conventional Dendritic Cells” in Advances in Immunology, F. W. Alt, Ed. (Academic Press, 2017), pp. 89–135.

26. M. L. Broz, Dissecting the Tumor Myeloid Compartment Reveals Rare Activating Antigen-Presenting Cells Critical for T Cell Immunity. Cancer Cell 26, 638–652 (2014).

27. Y. Lavin, Innate Immune Landscape in Early Lung Adenocarcinoma by Paired Single-Cell Analyses. Cell 169, 750–765 17 (2017).

28. A. R. Sánchez-Paulete, Cancer Immunotherapy with Immunomodulatory Anti-CD137 and Anti-PD-1 Monoclonal Antibodies Requires BATF3-Dependent Dendritic Cells. Cancer Discov 6, 71–79 (2016).

29. H. Salmon, Expansion and Activation of CD103+ Dendritic Cell Progenitors at the Tumor Site Enhances Tumor Responses to Therapeutic PD-L1 and BRAF Inhibition. Immunity 44, 924–938 (2016).

30. K. C. Barry, A natural killer–dendritic cell axis defines checkpoint therapy–responsive tumor microenvironments. Nat Med 24, 1178–1191 (2018).

31. D. G. Kirsch, A spatially and temporally restricted mouse model of soft tissue sarcoma. Nat. Med 13, 992–997 (2007).

32. A. T. Satpathy, Zbtb46 expression distinguishes classical dendritic cells and their committed progenitors from other immune lineages. J Exp Med 209, 1135–1152 (2012).

33. K. Kabashima, Intrinsic lymphotoxin-beta receptor requirement for homeostasis of lymphoid tissue dendritic cells. Immunity 22, 439–450 (2005).

34. W. Kc, L-Myc expression by dendritic cells is required for optimal T-cell priming. Nature 507, 243–247 (2014).

35. R. J. DeBerardinis, T. Cheng, Q’s next: the diverse functions of glutamine in metabolism, cell biology and cancer. Oncogene 29, 313–324 (2010).

36. L. Scourzic, E. Mouly, O. A. Bernard, TET proteins and the control of cytosine demethylation in cancer. Genome Med 7, **9** (2015).

37. L. Peng, A. Schousboe, L. Hertz, Utilization of alpha-ketoglutarate as a precursor for transmitter glutamate in cultured cerebellar granule cells. Neurochem Res 16, 29–34 (1991).

38. J. Chen, De novo nucleotide biosynthetic pathway and cancer. Genes & Diseases 10, 2331–2338 (2023).

39. K. E. Wellen, The hexosamine biosynthetic pathway couples growth factor-induced glutamine uptake to glucose metabolism. Genes Dev 24, 2784–2799 (2010).

40. T. Sathaliyawala, Mammalian target of rapamycin controls dendritic cell development downstream of Flt3 ligand signaling. Immunity 33, 597–606 (2010).

41. C. Guo, SLC38A2 and glutamine signalling in cDC1s dictate anti-tumour immunity. Nature (2023). 10.1038/s41586-023-06299-8.

42. C. Sinclair, mTOR regulates metabolic adaptation of APCs in the lung and controls the outcome of allergic inflammation. Science 357, 1014–1021 (2017).

43. P. Nicklin, Bidirectional transport of amino acids regulates mTOR and autophagy. Cell 136, 521–534 (2009).

44. J. L. Jewell, Differential regulation of mTORC1 by leucine and glutamine. Science 347, 194–198 (2015).

45. D. Meng, Glutamine and asparagine activate mTORC1 independently of Rag GTPases. J Biol Chem 295, 2890–2899 (2020).

46. M. Manifava, et al., Dynamics of mTORC1 activation in response to amino acids. Elife 5, e19960 (2016).

47. J. Encarnación-Rosado, et al., Targeting pancreatic cancer metabolic dependencies through glutamine antagonism. Nat Cancer 5, 85–99 (2024).

48. M. V. Recouvreux, et al., Glutamine mimicry suppresses tumor progression through asparagine metabolism in pancreatic ductal adenocarcinoma. Nat Cancer 5, 100–113 (2024).

49. S. Devalaraja, et al., Tumor-Derived Retinoic Acid Regulates Intratumoral Monocyte Differentiation to Promote Immune Suppression. Cell 180, 1098–1114.e16 (2020).

50. E. F. Garcia, The mitochondrial Ca2+ channel MCU is critical for tumor growth by supporting cell cycle progression and proliferation. Frontiers in Cell and Developmental Biology 11 (2023).

51. M. T. Dang, F. Mafra, M. Haldar, Isolation of myeloid cells from mouse brain tumors for single-cell RNA-seq analysis. STAR Protoc 2, 100957 (2021).

